# HERITABILITY AND QTL MAPPING OF AERIAL ROOTS AND OTHER YIELD COMPONENT TRAITS WITH IMPLICATIONS FOR N_2_ FIXATION IN *ZEA MAYS*

**DOI:** 10.1101/2025.09.10.675198

**Authors:** David A. O’Donnell, Jinliang Yang, Pablo Zamora, Anne Lorant, Chenyong Miao, Allen Van Deynze, Alan Bennett, Jeffrey Ross-Ibarra

## Abstract

*Zea mays* L. spp. *mays* (hereafter maize) populations from an indigenous community of Mexico have been reported to perform well using traditional cultivation practices that exclude industrial fertilizers despite low soil nitrate levels. The local maize cultivar is characterized by extended maturity, tall stature, and presence of thick aerial roots that secrete an abundance of polysaccharide-rich root exudate, or mucilage, that is implicated in the recruitment of diazotrophic microbes to facilitate biological nitrogen fixation. Here we estimate the broad sense heritability of traits related to nitrogen fixation in a panel of *Zea* entries spanning pre-domestication, post-domestication and post-improvement, and identify QTL via F2:F3 families for traits including aerial root node counts and various other yield related traits (*i.e.* germination rate, time to reproductive maturity, total biomass and nitrogen content of shoot and grain). Across two separate field studies, aerial root node count (AR) demonstrates heritability of 64% and 73%, and genetic mapping reveals three distinct QTL distributed on chromosomes 1 and 9. Further, we report novel QTL for germination rate via stand counts after direct sowing (SC), Plant Total Nitrogen (PTN), Plant Dry Mass (PDM), and a joint QTL for Grain Total Nitrogen (GTN) and Grain Dry Mass (GDM). Finally, we identify overlap between QTL for multiple traits (including AR and SC) and regions of elevated introgression from wild *Zea mays* L. spp. *mexicana* (hereafter *mexicana*) into Totontepec maize, with a greater degree of overlap than expected under a uniform genomic distribution suggesting adaptive introgression from *mexicana*. Given its pronounced aerial root morphology, we propose that *mexicana* is the ancestral source for prominent aerial roots and corresponding abundant mucilage production in Totontepec maize and other maize traditional varieties.

**ARTICLE SUMMARY:** *Zea mays* aerial roots (AR) and mucilage have been implicated in diazotrophic recruitment. This study estimates heritability and identifies QTL for AR and yield components via a diverse panel of *Zea* entries and F2:F3 mapping populations. Genetic mapping reveals three QTL for AR and several QTL for other yield traits, both novel and previously reported. This study also compares QTL and regions of elevated introgression from wild *Zea mays* L. spp. *mexicana* (hereafter *mexicana*) into a native cultivar. For AR QTL, enrichment for *mexicana*-derived haplotypes is greater than expected under a uniform genomic distribution, suggesting adaptive introgression from *mexicana*.

## INTRODUCTION

*Zea mays* L. spp. *mays* (hereafter maize), with a global output exceeding 48.6 billion bushels in the 2023–2024 market year, is the most widely produced crop in the world (World of Corn, 2024). Aside from traditional use for human consumption, maize has several other important applications (*i.e.* livestock feed, cooking oil, biofuel), and projections estimate that more than half of cereal crop calories will come from maize as the global human population continues to rise (Yan *et al*., 2011). The projected global human population of 9.75 billion by the year 2050 [under a medium population growth scenario] is expected to require 47% more crop calories relative to a 2011 baseline (Sands *et al*., 2023).

With the advent of the Green Revolution, historical increases in crop yield have been, at least in part, due to intense application of synthetic nitrogen fertilizers. Greater than 50% of such fertilizers are used for cereal production (Heffer, 2013), which doubled between 1966 and 2000 (Khush, 2001; Yan *et al*., 2011). Yet fertilizer use by cereals is inefficient, being utilized at a rate of less than 50% (Halvorson *et al*., 2002). This inefficiency results in nitrate leaching into soil and groundwater supplies, posing significant environmental concerns and health risks (Santi *et al*., 2013). Further, present rates of fertilizer over-application are leading to diminishing returns of crop yield via soil acidification and degradation (Robertson and Vitousek, 2009; Büyükkılıç Yanardağ, 2024), leaving an ever-growing need to develop alternative solutions for increased crop yield. This has stimulated research efforts to investigate the potential for biological nitrogen fixation (BNF) in cereal grains of agronomic importance (Christiansen-Weniger, 1998; Coque *et al*., 2006; Montañez *et al*., 2009; Montañez *et al*., 2012; Roesch *et al*., 2006; Santi *et al*., 2013; Van Deynze *et al*., 2018).

Diazotrophic bacterial communities that can fix atmospheric nitrogen are enriched in maize rhizospheres, but the addition of nitrogen fertilizer significantly reduces the abundance of these microbes (Roesch *et al*., 2006). Other researchers have demonstrated BNF in various maize cultivars using ^15^N isotope-dilution, and have successfully isolated diazotrophic bacteria cultures from endophytic maize tissues (Montañez *et al*., 2009). These workers also reported significant increases in shoot, root, and grain biomass for various maize cultivars in response to inoculation by diazotrophic bacteria, as well as positive correlations between biomass and % nitrogen derived from atmosphere (NDFA) as assessed via ^15^N isotopic dilution (Montañez *et al*., 2009).

Totontepec Villa de Morelos (hereafter Totontepec) is a small municipality in the state of Oaxaca, Mexico. Located in the Sierra Mixe region at over 1800 m elevation, the area has a mild climate (∼16.6 ± 3⁰C) and receives substantial rainfall (2063 mm annually), with peak precipitation occurring between June and October (Totontepec Villa de Morelos Climate, 2023). Traditional maize fields in this community are cultivated with minimal use of fertilizers or pesticides (Estrada *et al*., 2002), and several fields exhibit low soil nitrate levels. Notably, Totontepec maize performs well under such conditions and can reach heights exceeding 5 m at maturity (Van Deynze *et al*., 2018). This traditional variety is characterized by the presence of thick, abundant aerial roots that secrete significant amounts of polysaccharide-rich root mucilage, which has been shown to have a vital role to attract diazotrophic microbes and thereby facilitate BNF to benefit this cultivar (Van Deynze *et al*., 2018; Pankievicz *et al*., 2022). Van Deynze *et al*. (2018) first demonstrated this capacity using multiple approaches, including ^15^N natural abundance assays, acetylene reduction assays, and 16S ribosomal RNA metagenomic sequencing. This work revealed an enrichment of diazotrophic bacteria in mucilage derived from Tontontepec aerial root tissues, along with a low oxygen concentration gradient in aerial root mucilage to enable proper functionality of the nitrogenase enzyme in diazotrophic bacteria. Subsequent research has shown that BNF and mucilage production on Totontepec maize are governed by aerial root development and polysaccharide secretion from border cells, with thicker early-stage aerial roots (up to 9mm diameter, 0.5 cm length) produce the largest volume of mucilage (up to 2 mL per root) (Pankievicz *et al*., 2022). Moreover, in aerial roots exposed to water, RNA-seq analysis have revealed differential expression of genes involved in polysaccharide synthesis and degradation pathways, as well as ammonium and nitrate transporters. These findings support a functional link between mucilage secretion, diazotrophic microbial recruitment, and BNF in aerial root mucilage of this cultivar (Pankievicz *et al*., 2022).

Here, we aim to interpret trait heritability and characterize genetic loci associated with aerial root development and other yield component traits in *Zea mays*. Additional traits examined include germination rate, leaf tissue ^15^N abundance as an indicator of NDFA (Boddey, 1987), time to reproductive maturity, plant height, total biomass and nitrogen content of shoot and grain. To this end, we report phenotypic results from a genus *Zea* trial (hereafter *Zea* Trial 1), which includes Totontepec maize, multiple other traditional varieties from highland and lowland regions within and outside of Oaxaca, improved elite hybrid and inbred lines, and three wild teosinte taxa – *Zea mays* L. spp. *mexicana* (hereafter *mexicana*), *Z. mays* L. spp. *parviglumis* (hereafter *parviglumis*) and *Z. diploperennis* (hereafter *diploperennis*). As a complementary exploration, we identify the environmental conditions required to stimulate mucilage production from aerial roots under controlled greenhouse conditions. Further, we characterize traits within a nested association population of F2:F3 families derived from Totontepec maize and three elite inbred lines, and identify corresponding Quantitative Trait Loci (QTL) for aerial root abundance and other yield-related traits of interest.

Finally, we compare the identified QTL with genomic regions showing introgression from *mexicana* into Totontepec maize, and highlight introgression regions overlapping with QTL. *Mexicana* and *parviglumis* diverged from one-another approximately 30,000 years ago, while modern maize diverged from *parviglumis* roughly 12,000 years ago (Chen *et al*., 2022). Relative kinship between highland maize and *parviglumis* has been impacted by significant admixture between highland maize and *mexicana* (van Heerwaarden *et al*., 2010; Hufford *et al*., 2012; Hufford *et al*., 2013; Chen *et al*., 2022), which remain inter-fertile and overlap in geography (Matsuoka *et al*., 2002). Notably, *mexicana* shares overlapping geography with Totontepec maize and possesses prominent aerial roots with capacity for mucilage formation. Recent investigation of more than 1,000 wild and domesticated genomes shows that between 15 – 25% of the modern maize genome derives from introgression from *mexicana* (Yang *et al*., 2023). We hypothesize that maize aerial roots and related yield-component traits are genetically complex, governed by multiple interacting loci (similar to other pre-domestication traits (Buckler *et al*., 2009; Tian *et al*., 2011)) and with key allelic contributions derived from *mexicana* via introgression into Totontepec maize. Ultimately, identifying the genetic basis of traits relevant to BNF and efficient yield gains in maize may support selective breeding for more efficient maize varieties that require significantly less synthetic fertilizer, thereby reducing both production costs and environmental impacts.

## MATERIALS AND METHODS

### Plant Materials, Growth Conditions and Experimental Design

#### *Zea* Trial 1

*Zea* entries are listed in Table S1, with corresponding map positions shown in Figure S1. Entries include lowland and highland traditional variety populations within and outside of Oaxaca, wild teosintes (*mexicana*, *parviglumis*, and *diploperennis*), and improved conventional groups, including three inbred lines and one hybrid. Three subpopulations of Totontepec maize from different farmers are distinguished by field location and designated as Totontepec cultivars 1, 2, and 3. We performed seed germinations with heavy-duty brown germination paper in a sterilized 26⁰C growth chamber with a 16 hour light: 8 hour dark cycle. After 12 days, we transplanted seedlings to 48-well planter trays with a 1:1 mixture of perlite: vermiculite. To ensure plant vigor during early growth stages, we maintained plants in a greenhouse for 18 days with daily irrigation prior to transplantation to field conditions. We supplemented irrigation with fertilizer according to UC Davis Core Greenhouse Facility standards (UC Davis Research Greenhouses, c2009). We transplanted plants to the field in a complete randomized block design with 50 cm spacing between adjacent plants. To simulate rainfall characteristic of the native Oaxacan environment, we cultivated plants within a mist irrigation system; during vegetative and reproductive growth phases, we applied mist treatment for approximately 2 hours per day to stimulate root exudate production from aerial roots.

#### Greenhouse Aerial Root Mucilage Stimulation

We assessed four subgroups of *Z. mays*, including Totontepec maize, *mexicana* and *parviglumis*, and one improved conventional hybrid ‘B73 x Mo17’. Totontepec maize seeds were derived from a single plant of Totontepec cultivar 2, while *mexicana*, *parviglumis* and ‘B73 x Mo17’ seeds each were derived from a single accession from the USDA genebank (respectively, PI 566674, Ames 21809, Ames 19097). We grew plants within two soil-floor greenhouse chambers in a randomized complete block design of identical layout per chamber, with 21 seeds sown per entry per chamber (direct sowing) and variable final germination. We did not treat soil with fertilizer prior to the experiment. In chamber 1, we utilized solely drip irrigation to sustain plant growth and development. In chamber 2, we utilized misting derived from an overhead system to simulate rainfall, supplemented with drip-irrigation to maintain consistent overall water quantity applied per treatment. We applied mist periodically to maintain humidity levels ≥80%.

#### *Zea* Trial 2

We made three F2:F3 mapping populations in a nested association design, derived from single F1 plants resulting from the cross between Totontepec cultivar 2 as female parent and three inbred testers – Mo17 (PI 558532), LH82 (PI 601170) and B73 (PI 550473). After selfing F2s, we obtained 77, 119, and 59 F_2_:F_3_ families for bi-parental populations derived from B73, LH82, and Mo17, respectively. We planted F_2_:F_3_ families within field conditions in randomized row design, with two replicates per family for most lines. Each replicate included 15 seeds at 8-inch spacing in single row format (direct sowing to field), with commercial maize border rows between each entry row. We attempted field enrichment with 0.7% ^15^N distributed over two applications (pre-planting and at week 5 post-germination). We applied minimum nitrogen fertilizer at 80 kg/ha, also distributed over two applications (pre-planting and week 5 post-germination, 40 kg/ha per application). We furrow-irrigated plants for the initial 7 weeks of development. At week 8, we began to apply sprinkle irrigation for approximately one hour per day to simulate regular rainfall; this continued for the remainder of the experiment. At week 11 post-germination, we secured large plants with a trellis system.

### Soil Analyses

#### *Zea* Trial 1

We sampled soil from fifteen distinct locations prior to the experiment, and from six distinct locations upon conclusion of the experiment, evenly distributed throughout the field and at four depths per location (5 cm, 25 cm, 50 cm, 100 cm). We oven-dried samples at 50 °C for 24 hours and analyzed for nitrogen (NO_3_-N), phosphorus (Olsen-P), and potassium (Exchangeable-K) content as determinants of soil fertility, following the specifications detailed by the UC Davis Analytical Lab (UC Davis Analytical Laboratory, c2000–2014). We analyzed samples for ^15^N natural abundance as specified by the UC Davis Stable Isotope Facility (UC Davis Stable Isotope Facility, c2020). We averaged nutrient and ^15^N values across field locations according to depth and time of sampling; values are shown in Figure S2.

#### *Zea* Trial 2

We sampled soil from three locations prior to the experiment at three depths per location (5 cm, 25 cm, 50 cm) and from nine locations upon conclusion of the experiment (at four depths of 5 cm, 25 cm, 50 cm, 75 cm per location), evenly distributed throughout the field. We oven-dried samples at 50 °C for 24 hours and analyzed for nitrogen (NO_3_-N), phosphorus (Olsen-P), and potassium (Exchangeable-K) content as determinants of soil fertility, according to the specifications detailed by the UC Davis Analytical Lab (UC Davis Analytical Laboratory, c2000–2014). We analyzed samples for ^15^N content as specified by the UC Davis Stable Isotope Facility (UC Davis Stable Isotope Facility, c2020). We averaged nutrient and ^15^N values across field locations according to depth and time of sampling; such values are displayed in Figure S5.

### Phenotyping and Statistical Analysis

#### *Zea* Trial 1

We recorded stand counts after direct sowing (SC) for each plot at 2 weeks after transplant. For ^15^N natural abundance assessment (Boddey, 1987; Boddey *et al*., 2001; Bremer and Van Kessel, 1990), we sampled plant leaf tissue at two separate time-points of 9 and 13 weeks post-transplant to the field (^15^N_9W_, ^15^N_13W_). Improved varieties (including B73, Mo17, Hickory King, and the B73 x Mo17 F1 hybrid) flowered shortly after time-point 1, and thus were not sampled for time-point 2. We sampled ∼100 mg of leaf tissue from the third-youngest leaf of each plant, dried tissue at 50 °C for 24 hours, and analyzed it for ^15^N content as above. Theoretical values of NDFA are expressed as a percentage of total plant nitrogen, based on comparison to ‘reference’ negative control conventional B73 (an improved inbred that demonstrates a ^15^N signature indicative of minimal/no association to diazotrophic bacteria). We calculated NDFA values using the equation %NDFA = ((^15^N_reference_ – ^15^N_experimental_) / (^15^N_reference_)) * 100 (Bremer and Van Kessel, 1990). We also surveyed plants once per week to assess days to tassel emergence (DT) and pollen shed (DPS). At maturity, we measured plants for total height (above-ground shoot length from soil to tip of tassel) and number of nodes possessing aerial roots (AR). We performed Analysis of Variance (ANOVA) and estimated broad-sense heritability using R version 3.3.1 GUI 1.68 Mavericks build. We generated bar charts using Microsoft Excel version 15.32.

#### *Zea* Trial 2

As an indication of germination rate, we recorded SC for each plot at 3 weeks after direct sowing to open field. For ^15^N natural abundance assessment, we sampled plants for three time-points at weeks 8, 10, and 12 post-germination (^15^N_8W_, ^15^N_10W_, ^15^N_12W_, respectively). Cumulatively, we sampled ∼100 mg of leaf tissue from the third-youngest leaf for ∼5 plants per F_2_:F_3_ family, dried at 50 °C for 24 hours, and analyzed for ^15^N content. We surveyed families once per week to assess DT and DPS. At maturity, we scored three plants per F_2_:F_3_ family for total number of AR nodes, total plant dry mass above the soil interface excluding grain (PDM), and total grain dry mass (GDM). We bulked dry plant biomass per family (one bulk of plant shoot and one bulk of grain), finely ground and further processed for plant total nitrogen (PTN), plant total nitrogen percentage (PTNP), grain total nitrogen (GTN) and grain total nitrogen percentage (GTNP) (UC Davis Analytical Laboratory, c2000–2014). We performed ANOVA and estimated broad-sense heritability using R version 3.3.1 GUI 1.68 Mavericks build. We adjusted scores for phenotypes of interest using Best Linear Unbiased Prediction (BLUP) with the ‘lme4’ package (Bates *et al*., 2015) in RStudio v.1.0.136 (R Studio Team, 2022), to account for environmental effects or uneven distribution of ^15^N applications. Based on leaf tissue sample analysis for ^15^N, we determined that, out of 61 total beds, field ^15^N enrichment application was disproportionately elevated in beds 3, 5, 7, 9 and 11. For plants derived from such beds, we excluded ^15^N values from further phenotypic data analysis and genetic mapping efforts. For BLUP adjustment of collected phenotype data, we used replicate number, field location coordinates (bed and row number), and biparental subpopulation as fixed effects, and genotype as random effect in the model. We generated all histograms and statistical outputs via TASSEL v.5 software (Bradbury *et al*., 2007).

### Plant Genotyping and QTL Mapping

#### *Zea* Trial 2

For parental lines and each F_2_:F_3_ family, we collected a total amount ∼100 mg of leaf tissue per plot (bulk sampling). We extracted DNA from each sample using the Qiagen DNeasy Plant Mini Kit (DNeasy Plant Mini Kit, c2013–2017). We genotyped samples via Genotyping-by-Sequencing (GBS), as previously described by Elshire *et al*. (2011). Single nucleotide polymorphism (SNP) calling was performed against the B73 reference genome version 2 (RefGen_v2 / AGPv2; GCA_000005005.4) from the NCBI RefSeq database (Goldfarb *et al*., 2025). This yielded 955,690 SNPs, and filtration with a call rate ≥70% reduced the marker set to 30,757 markers. To obtain a composite genotype for the Totontepec cultivar parent (not a fixed line), we compared genotypes of eight bulk samples from eight Totontepec plots to identify all loci fixed across samples (including all loci with available data for at least 7 sample bulks). The composite Totontepec genotype included 8,347 SNP markers fixed across Totontepec sample bulks. Using this set of markers, pairwise genotype comparison of Totontepec to each inbred parent yielded 1,033; 1,673; and 1,064 informative markers respectively for bi-parental populations derived from B73, LH82 and Mo17. We used the Genotype Corrector method (Miao *et al*., 2018) to correct for false recombinants caused by genotyping errors in GBS data. For subsequent QTL mapping steps noted herein, we used the ‘qtl’ package (Broman *et al*., 2003) in RStudio v.1.0.136 (R Studio Team, 2022). Per bi-parental population, we organized linkage groups with max.rf=0.35 and min.lod=9. We determined LOD (Logarithm of the Odds) thresholds via permutation testing with 1000 permutations, and thereafter performed composite interval mapping per trait using BLUP phenotype values via the Kosambi mapping function.

### Admixture Analysis of *mexicana* into Totontepec Maize

To investigate introgression from *mexicana* into Totontepec maize, we used genotyping data from n=94 maize landraces, n=130 *parviglumis* accessions, and n=120 *mexicana* accessions (Hufford *et al*., 2013) as reference populations. Admixed chromosomes were modeled as mosaics of ancestral segments originating from different reference populations, and ancestry was inferred using HapMix (Price *et al*., 2009), with haplotypes for reference populations pre-phased using fastPHASE (Scheet *et al*., 2006). For each chromosome, a jackknife resampling strategy was applied to assess the robustness of local ancestry estimates. The resulting genome-wide matrix was averaged across replicates to generate a consensus estimate of *mexicana* ancestry. Thereafter, we compared regions of elevated *mexicana* introgression (defined by *mexicana* haplotype frequency >50% for a given region) against QTL detected herein to identify regions of common overlap.

## RESULTS

In *Zea* Trial 1, traits of interest show clear phenotypic variation both within and among species. Figure 3 illustrates variation in NDFA and AR traits across groups of *Zea*, with detailed distributions shown in Figures S3 and S4a. NDFA varied substantially across populations, but was relatively stable within populations and across time-points (Figure S3). Broad-sense heritability estimates for NDFA were 67.2% at 9 weeks and 60.7% at 13 weeks (Table 1), indicating a strong genetic component.

**Table 1:** Broad-Sense Heritability for Traits of Interest. Traits are listed at far left. Columns 2 – 3 indicate broad-sense heritability estimates from *Zea* Trial 1 and *Zea* Trial 2, respectively. For *Zea* Trial 2, heritability estimates were calculated for the population excluding parental groups (to include only F_2_:F_3_ families). Key: NDFA = Nitrogen Derived from Atmosphere at key time-points per trial; SC = stand count after direct sowing to field; DPS = ∼ days to pollen shed; AR = aerial root nodes. PH = plant height at maturity from plant base to tip of tassel; PDM = mature plant total dry mass; PTN = mature plant total nitrogen content; PTNP = mature plant total nitrogen content percentage; GDM = mature grain total dry mass; GTN = mature grain total nitrogen content; GTNP = mature grain total nitrogen content percentage.

| Trait | Heritability<br>(*Zea* Trial 1) | Heritability<br>(*Zea* Trial 2) |
| --- | --- | --- |
| NDFA<sub>9w</sub> (9 weeks) | 0.672 | - |
| NDFA<sub>13w</sub> (13 weeks) | 0.607 | - |
| NDFA<sub>8w</sub> (8 weeks) | - | 0.231 |
| NDFA<sub>10w</sub> (10 weeks) | - | 0.189 |
| NDFA<sub>12w</sub> (12 weeks) | - | 0.092 |
| SC | - | 0.780 |
| DPS | 0.930 | 0.923 |
| AR | 0.640 | 0.730 |
| PH | 0.880 | - |
| PDM | - | 0.398 |
| PTN | - | 0.514 |
| PTNP | - | 0.329 |
| GDM | - | 0.450 |
| GTN | - | 0.429 |
| GTNP | - | 0.361 |

Additional traits, including AR abundance (Figure 3, Figure S4a), plant height (Figure S4b), and time to maturity (days to tassel [DT] and days to pollen [DPS]; Figure S4c), also showed significant inter-population variation and high heritability: 64.0% for AR, 88.0% for height, and 93.0% for maturity traits (Table 1).

Aerial root (AR) morphology and abundance varied notably among wild teosintes. *Mexicana* displays prominent aerial roots (Figure 1, 2), whereas *parviglumis* (Figure 2) and *diploperennis* (not shown) display smaller structures. *Diploperennis* produced more aerial roots on average (5.29 ± 1.38 s.d. per plant) than *mexicana* (3.47 ± 1.91 s.d. per plant), but these were smaller and did not produce mucilage (Figure S4a). Both *mexicana* and *diploperennis* had more AR-bearing nodes than *parviglumis* (Figure S4a).

**Figure 1:**
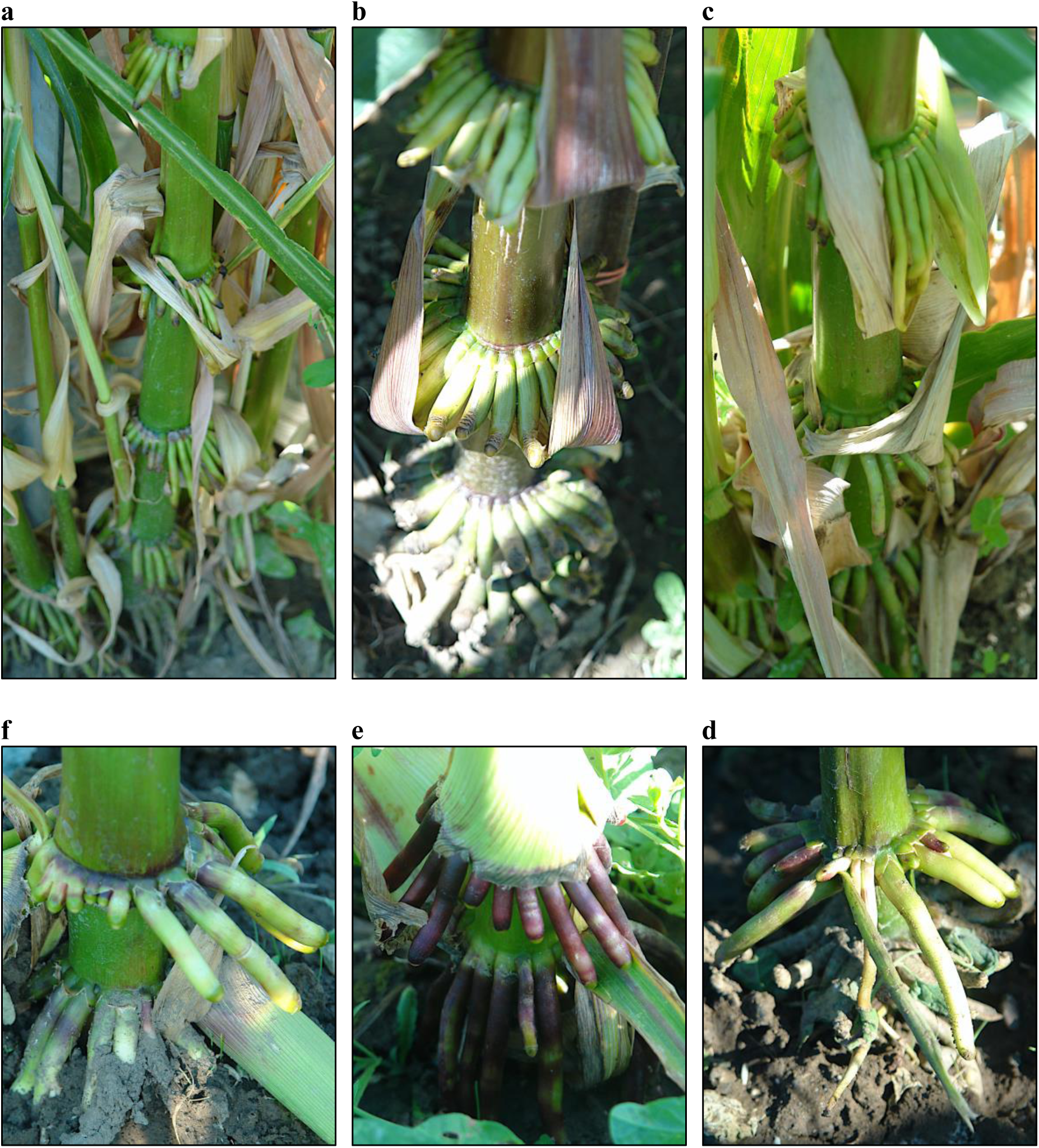
Aerial Root Morphology – *Mexicana*, Totontepec and Improved Entries. Photos of aerial root morphologies at reproductive maturity for the following entries (clockwise from top-left): (**a**) teosinte *mexicana*, (**b**) Totontepec cultivar 2, **(c)** Totontepec cultivar 3, (**d**) improved hybrid B73xMo17, (**e**) improved inbred Mo17, and (**f**) improved inbred B73.

**Figure 2:**
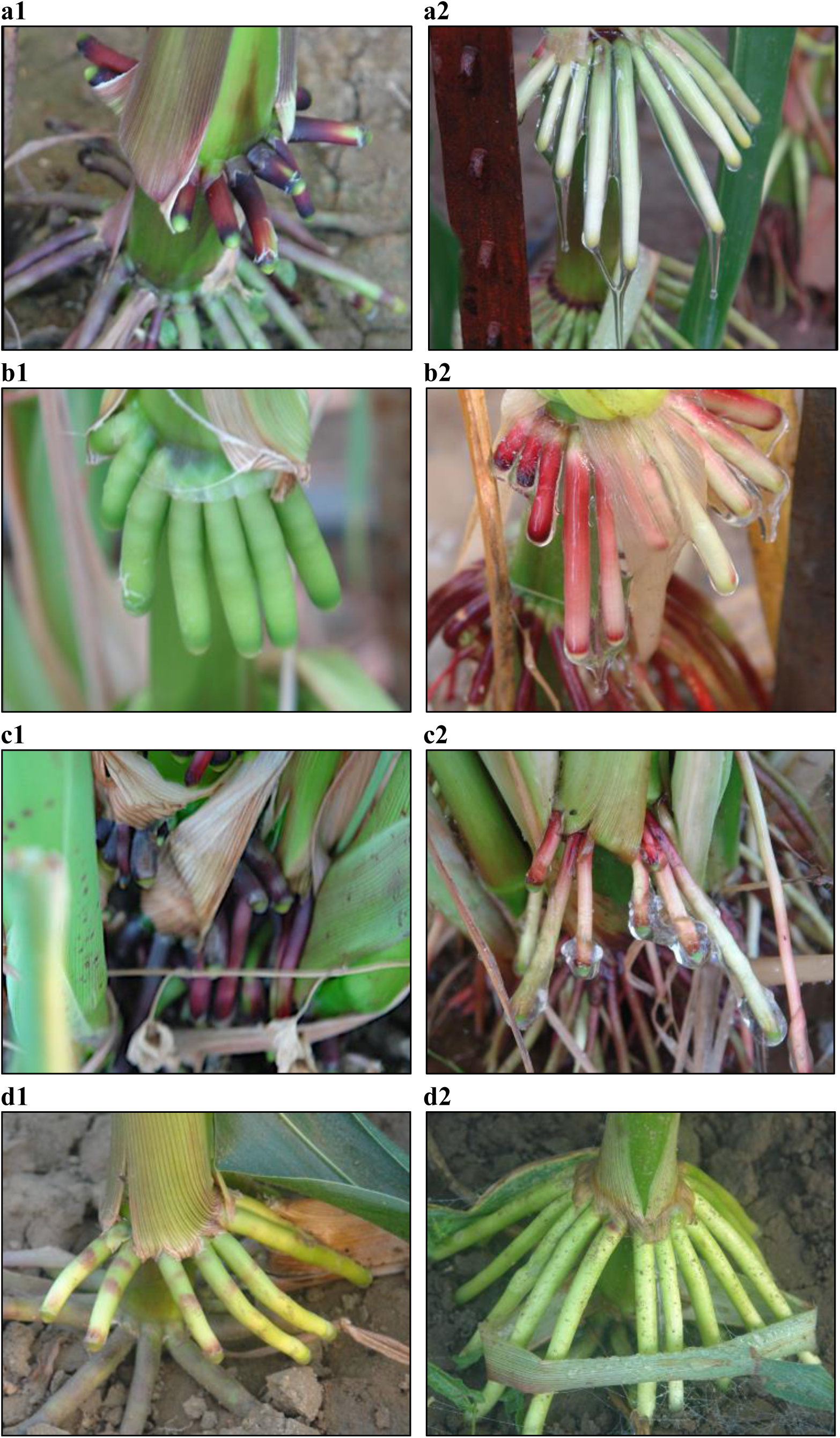
Aerial Roots in Drip-Irrigated *vs.* Misted Greenhouse Conditions. Series of photographs showing aerial roots for *Zea mays* subgroups within misted and drip-irrigated conditions. From top to bottom: (**a**) Totontepec cultivar 2, (**b**) teosinte *mexicana*, (**c**) teosinte *parviglumis*, and (**d**) improved hybrid ‘B73 x Mo17’, respectively. Left panels (**1**) = drip-irrigated conditions; right panels (**2**) = mist conditions. All photos taken 66 days after sowing of seeds.

To evaluate capacity for mucilage production across different groups of *Zea*, we tested aerial root responses in a control treatment of low external humidity as well as a misting treatment to maintain 80% relative humidity (see Methods). As shown in Figure 2, mucilage was abundant in Totontepec maize, *mexicana* and *parviglumis* under misted conditions but absent in the control treatment. In contrast, B73 x Mo17 hybrid lacked mucilage in both treatments, likely due to minimal aerial root development, with most AR nodes proximal to or embedded in the soil, functioning as brace roots for plant stability.

Based on heritability estimates obtained in *Zea* Trial 1, we conducted QTL mapping in a nested F2:F3 population (*Zea* Trial 2), analyzing each of the three bi-parental subpopulations separately. Heritability estimates across both trials are summarized in Table 1. NDFA exhibited low heritability in *Zea* Trial 2, with the highest estimate of 23.1% at 8 weeks, likely reflecting the narrower genetic base of *Zea* Trial 2. In contrast, DPS (days to pollen shed) showed strong heritability at 92.3%. Distributions of DT, DPS, PDM, PTN and AR were approximately normal (Figure S6), with heritability estimates of 39.8%, 51.4%, and 73.0%, for PDM, PTN, and AR respectively (Table 1).

Initial QTL mapping of Zea Trial 2 revealed artificially inflated genetic map lengths, likely due to false recombination events arising from errors and missing data typical of GBS. Genotype correction following Miao *et al*. (2018) reduced map inflation and improved QTL detection (Figure S7). Across the three bi-parental groups, we identified 14 unique QTL across eight traits, each explaining 14.81% to 44.13% of phenotypic variance (Table 2, Figure 4). Traits related to maturity time (DT, DPS) were governed by the same QTL, and thus were consolidated as ‘DPS’ QTL. Consistent with low heritability estimates, no QTL were detected for NDFA.

**Figure 3:**
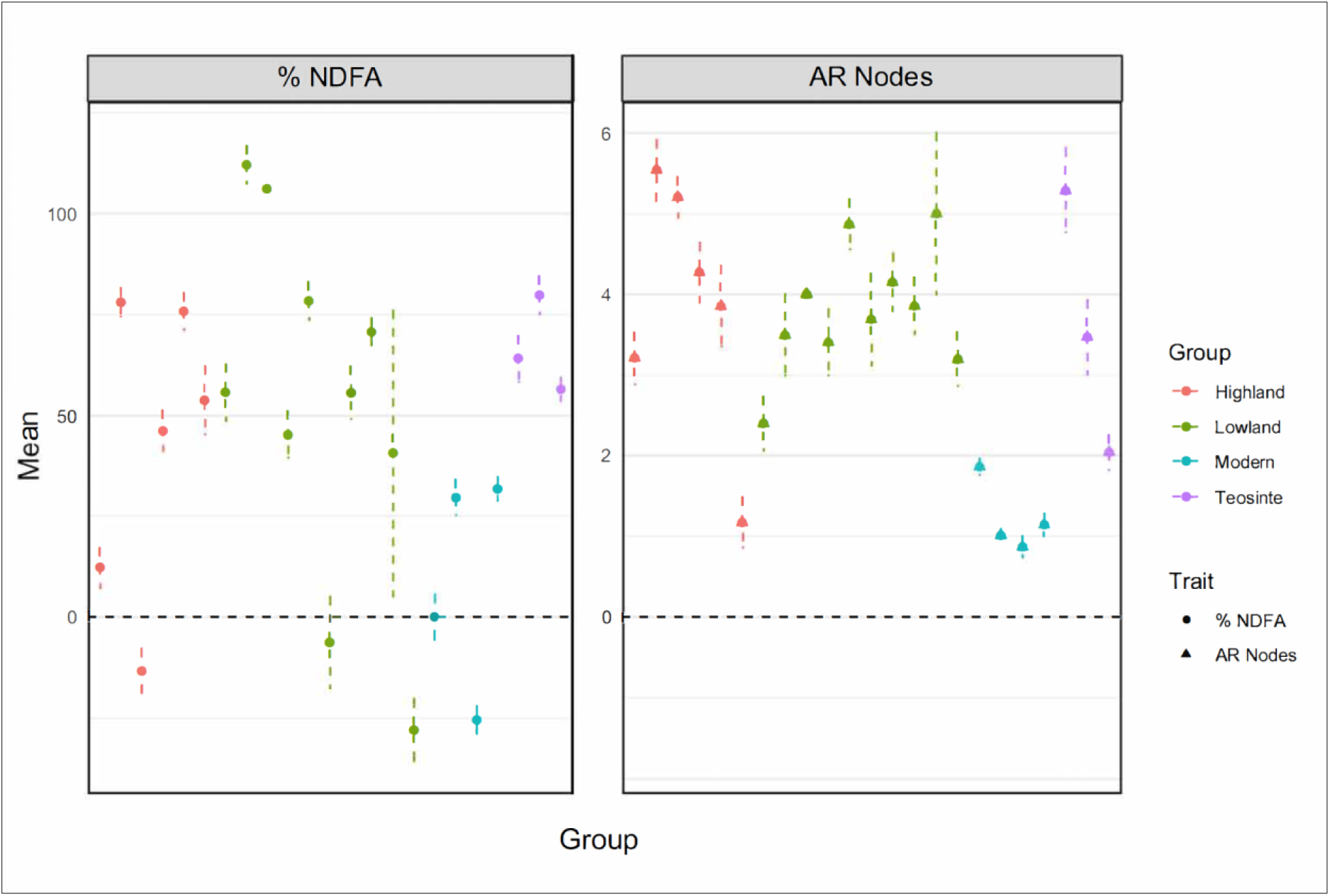
%NDFA and Aerial Root Node Variation among *Zea* Subgroups. For *Zea* Trial 1, plots displaying plot mean values for % NDFA (left plot) and aerial root node count (right plot) in fully mature plants of the different *Zea* groups (listed in Table S1), with standard error indicated by vertical dotted lines. Groups are color coded according to legend at right. For phenotype values corresponding to each entry, refer to Figure S3 and Figure S4a.

**Figure 4:**
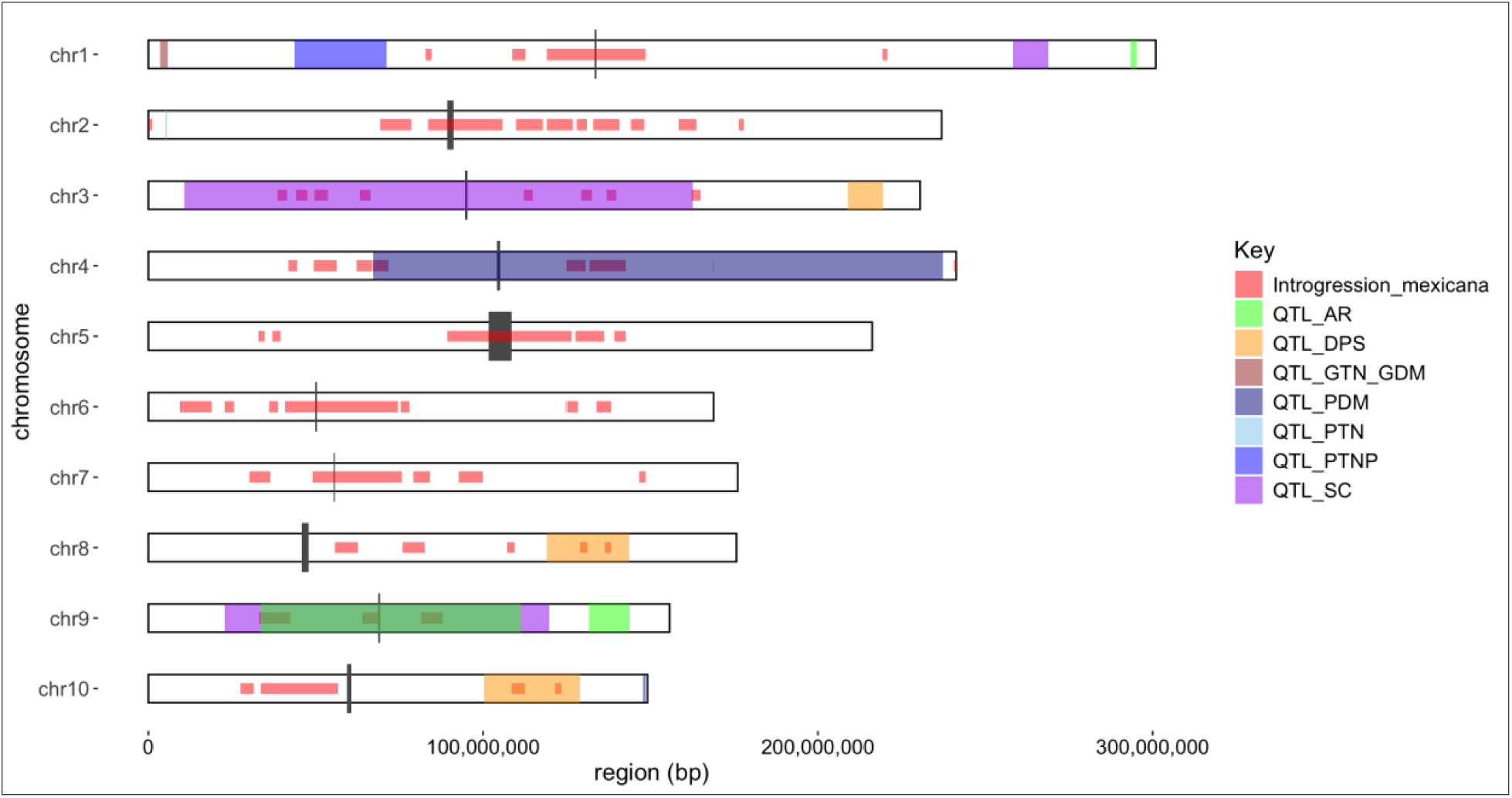
QTL and Elevated *Mexicana* Introgression into Totontepec Maize. Left Y-axis indicates chromosome number, with chromosomes shown in ascending order from top to bottom. Identified QTL per trait of interest, as well as regions of elevated introgression from *mexicana* into Totontepec cultivar (>0.5), are indicated by color along corresponding chromosome regions, with color key per trait at right side. Color Key: Introgression_mexicana = regions of elevated introgression from *mexicana* into Totontepec cultivar (>0.5); QTL_AR = QTL for ‘Aerial Root Nodes’; QTL_DPS = QTL for ‘Days to Pollen Shed’; QTL_GTN_GDM = QTL for ‘Grain Total Nitrogen’ and ‘Grain Dry Mass’ (overlapping QTL for highly related traits); QTL_PDM = QTL for ‘Plant Dry Mass’; QTL_PTN = QTL for ‘Plant Total Nitrogen’; QTL_PTNP = QTL for ‘Plant Total Nitrogen Percentage’; QTL_SC = QTL for ‘Stand Counts after direct sowing’. Shown alongside each QTL is its corresponding peak LOD score and % trait variance governed by QTL. See also Table 2.

**Table 2:** QTL, Confidence Interval Coordinates and %. Trait Variance List of QTL detected via composite interval mapping, for all traits across three bi-parental populations of *Zea* Trial 2 (Totontepec x: B73, LH82, Mo17). The table shows 1.5-LOD confidence intervals, peak scores per QTL, and % trait variance. The table is sorted first by trait of interest, with traits listed in column 1. Related traits are grouped together for ease of visualization. The population corresponding to each identified QTL is indicated by the elite inbred parent listed in column 2. Within trait categories, QTL are further sorted by chromosome position. Column 3 – 13, respectively, indicate: chromosome number, QTL name, interval start position (base pair coordinate), interval peak position (bp coordinate), interval end position (bp coordinate), interval start position (centiMorgans), interval peak position (cM), interval end position (cM), QTL peak score, physical length of significance interval (Megabase pairs), and genetic map length of significance interval (cM). Columns 14 – 16, respectively, indicate % trait variance governed by QTL, chi^2^ p-value, and p-value of the F-statistic. Column 17 indicates either novel QTL identified herein or else coincidence with QTL from prior literature, indicated by publication reference number as listed in bibliography.

| Trait | Inbred Parent | Chr | QTL Name | LOD Int. Start (bp) | LOD Int. Peak (bp) | LOD Int. End (bp) | LOD Int. Start (cM) | LOD Int. Peak (cM) | LOD Int. End (cM) | LOD Int. Peak Score | LOD Int. Total (Mb) | LOD Int. Total (cM) | % Var. | P value (Chi2) | P value (F) | Coincident QTL from Literature |
| --- | --- | --- | --- | --- | --- | --- | --- | --- | --- | --- | --- | --- | --- | --- | --- | --- |
| AR | LH82 | 1 | 1.4 | 293329598 | 294813085 | 295309095 | 246.94 | 251.50 | 271.86 | 6.25 | 1.98 | 24.92 | 18.44 | 0.00 | 1.04E-06 | Novel QTL |
| AR | LH82 | 9 | 9.2 | 33670979 | 102497421 | 111431608 | 91.97 | 97.10 | 103.21 | 4.27 | 77.76 | 11.24 | 18.89 | 0.00 | 7.71E-07 | Novel QTL |
| AR | MO17 | 9 | 9.3 | 131569767 | 139410267 | 143804017 | 117.69 | 122.30 | 127.10 | 4.20 | 12.23 | 9.41 | 21.61 | 0.00 | 4.29E-04 | (Zhang *et al.* 2018) |
| DPS | MO17 | 3 | 3.2 | 209024681 | 211685092 | 219411033 | 102.15 | 120.70 | 171.91 | 4.35 | 10.39 | 69.76 | 18.06 | 0.00 | 7.92E-05 | (Li *et al.* 2016) |
| DPS | MO17 | 8 | 8.1 | 118971709 | 123844193 | 143492696 | 152.81 | 155.50 | 159.10 | 6.23 | 24.52 | 6.29 | 44.13 | 0.00 | 5.75E-09 | (Du *et al.* 2021, Leng *et al.* 2022, Li *et al.* 2016) |
| DPS | B73 | 10 | 10.1 | 100252518 | 108117991 | 128967696 | 101.96 | 104.05 | 112.45 | 5.50 | 28.72 | 10.50 | 28.00 | 3.21E-06 | 5.26E-06 | (Du *et al.* 2021, Li *et al.* 2016, Zhang *et al.* 2018) |
| GDM | MO17 | 1 | 1.1 | 3408430 | 4996244 | 5813154 | 0.00 | 26.26 | 42.45 | 4.32 | 2.40 | 42.45 | 32.80 | 4.84E-05 | 8.79E-05 | Novel QTL |
| GTN | MO17 | 1 | 1.1 | 3408430 | 4996244 | 5813154 | 0.00 | 26.26 | 42.45 | 4.04 | 2.40 | 42.45 | 31.26 | 8.50E-05 | 1.49E-04 | Novel QTL |
| PTNP | MO17 | 1 | 1.2 | 43756650 | 61021047 | 71127215 | 115.52 | 121.81 | 124.51 | 4.44 | 27.37 | 9.00 | 29.72 | 3.61E-05 | 6.13E-05 | (Mandolino *et al.* 2018) |
| PTN | B73 | 2 | 2.1 | 4990077 | 5106835 | 5565686 | 71.19 | 89.28 | 92.09 | 6.30 | 0.58 | 20.89 | 32.32 | 2.97E-07 | 5.33E-07 | Novel QTL |
| PDM | MO17 | 4 | 4.1 | 67143858 | 237313637 | 237313637 | 37.22 | 233.98 | 233.98 | 3.91 | 170.17 | 196.76 | 19.00 | 2.22E-03 | 3.05E-03 | Novel QTL |
| PDM | LH82 | 10 | 10.2 | 147811484 | 147956241 | 148581023 | 162.68 | 184.76 | 198.66 | 4.07 | 0.77 | 35.97 | 14.80 | 8.50E-05 | 1.08E-04 | Novel QTL |
| SC | LH82 | 1 | 1.3 | 258353240 | 262937140 | 268813141 | 206.90 | 213.20 | 218.47 | 5.04 | 10.46 | 11.57 | 17.86 | 9.09E-06 | 1.22E-05 | Novel QTL |
| SC | B73 | 3 | 3.1 | 10683505 | 16566244 | 162587460 | 123.28 | 148.57 | 176.98 | 3.88 | 151.90 | 53.71 | 17.06 | 7.46E-04 | 9.88E-04 | (Han *et al.* 2014) |
| SC | MO17 | 9 | 9.1 | 22782253 | 108296473 | 119839322 | 100.05 | 110.67 | 114.35 | 5.65 | 97.06 | 14.29 | 15.03 | 8.88E-03 | 1.13E-02 | Novel QTL |

Given its importance to mucilage production and BNF, QTL associated with aerial root abundance were of particular interest. We detected QTLs on chromosome 1 (1.98 Mb, 18.44% trait variance) and 9 (77.76 Mb, 18.89% trait variance) in the LH82 bi-parental population, and an additional QTL on chromosome 9 (12.23 Mb, 21.61% trait variance) in the Mo17 background (Table 2, Figure 4). Confidence intervals for the two QTL on chromosome 9 did not overlap, suggesting that these QTL reflect distinct loci rather than a shared underlying signal.

We also explored introgression from *mexicana* into Totontepec maize (Figure 4, 5). Regions of elevated introgression, defined as genomic intervals with >50% *mexicana* haplotype frequency, were compared with QTL identified in *Zea* Trial 2. According to the B73 Ref_Gen_v2 genome annotation (Portwood *et al*., 2019), the maize genome contains 109,871 annotated genes, while there were 15,003 unique gene models within elevated introgression regions – ∼13.66% of the genome. Under a null hypothesis of random distribution, we would expect similar overlap of introgression with QTL regions. We observed 4,396 genes in AR QTL, with 774 genes (17.61%) also in introgression regions (Figure 4, Table S2, Table S3), indicating significant enrichment for *mexicana*-derived haplotypes within AR QTL regions (one-tailed chi-squared test, *P* <0.0001). We highlight a caveat that AR QTL 1.4 and 9.3 show no overlap with introgression segments, while AR QTL 9.2 shows correspondingly elevated enrichment; of 3,342 genes within AR QTL 9.2, 774 genes (23.16%) are in regions of elevated *mexicana* ancestry (Figure 4, Table S2, Table S3). This same region of introgression also overlapped with 18.63% of SC QTL 9.1, which is associated with germination rate and seeding vigor (Figure 4, Table S2, Table S3).

**Figure 5:**
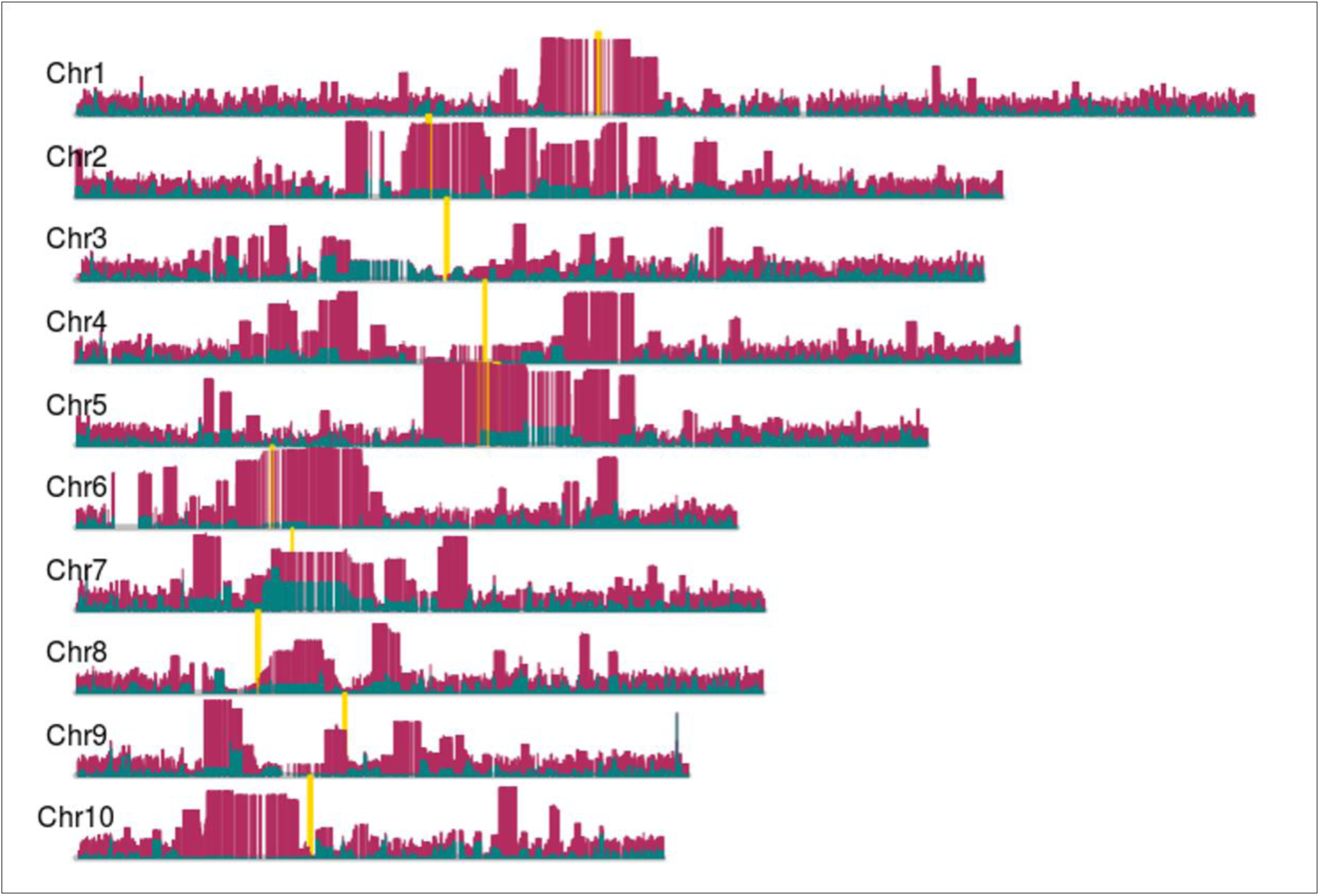
Genome-wide Landscape of Introgression from *Mexicana* into Totontepec Maize Lines. The x-axis represents the physical positions of the maize chromosomes (Chr1–Chr10), while the y-axis indicates the average probability of mexicana ancestry (magenta, the average estimate of *mexicana* introgression; teal, jackknife replicates). Yellow boxes indicate centromeric regions.

Additional overlaps were observed between *mexicana* introgression regions and QTL associated with other traits, including DPS, PDM, and SC (Figure 4, Table S2, Table S3). To determine whether these overlaps exceeded random expectations, we performed a genome-wide enrichment analysis. Per above, under a null hypothesis of random distribution, we would expect ∼13.66% of QTL regions to overlap with elevated signals of introgression. Respectively for DPS, PDM, and SC, we observed 14.13%, 9.46%, and 15.41% enrichment for *mexicana*-derived haplotypes (Table S3). Of these three traits, only SC QTL were significantly enriched beyond random expectation (one-tailed chi-squared test, *P* <0.0001).

## DISCUSSION

### Variation of Aerial Roots and Other Yield Component Traits throughout Genus Zea

Previous studies have demonstrated that Totontepec maize is capable of recruiting diazotrophic microorganisms via mucilage secreted from aerial roots and contributing to biological nitrogen fixation (BNF) as evidenced by nitrogen isotope signatures in field-grown plants (Van Deynze *et al*., 2018; Pankievicz *et al*., 2022). In the present study, we expanded upon these findings by characterizing aerial root nodes (AR), NDFA, and other yield component traits across a diverse panel of *Zea* accessions. *Zea* Trial 1 included wild teosintes (*parviglumis*, *mexicana,* and *diploperennis*), traditional maize varieties from highland and lowland regions (including three distinct Totontepec subpopulations), and improved materials comprising three elite inbred lines and one commercial hybrid (Table S1). This structured sampling allowed us to compare trait distributions across major evolutionary stages in maize, from wild progenitors to modern cultivars. *Zea* Trial 1 also served as the basis for estimating broad-sense heritability for a subset of traits. As expected, we observed high heritability for flowering time (DPS; H² = 93%) and plant height (H² = 88%) (Table 1), consistent with prior reports (Buckler *et al*., 2009; Peiffer *et al*., 2014). In contrast, AR and NDFA traits exhibited moderate heritability (60 – 68%; Table 1), indicating more complex genetic and environmental control. Importantly, we did not observe a clear, stepwise reduction of these traits across the evolutionary groups. For example, wild teosintes and traditional maize varieties exhibited broad and overlapping distributions for both AR node count and NFDA values. This suggests that these traits were not subject to strong directional selection during initial domestication from teosinte. However, we observed a marked reduction in both AR node number and NDFA signal in post-domestication entries (Figure 3, Figure S3, Figure S4a), suggestive of a degree of post-domestication trait erosion and potentially linked to modern breeding programs’ priorities on structural stability (such as brace root anchorage), uniform growth, and maximized yield under high-input systems. These factors may have inadvertently contributed to the loss or suppression of traits associated with aerial root mucilage secretion and symbiosis with diazotrophic microorganisms.

Improved maize lines typically present only one to two above-ground root-bearing nodes per plant. Upon soil contact, these roots transition to brace roots, contributing primarily to mechanical stability. The relative absence of mucilage-producing aerial roots in these improved lines (Figures 1, 2 and 3), along with their reduced NDFA signatures (Figure 3), suggests a mechanistic link between aerial root presence and NDFA. However, this relationship is not consistent, as some traditional varieties with prominent AR still displayed low NDFA (Figures S3 and S4), and NDFA varied substantially among Totontepec subpopulations with similar AR morphologies (Figures S3 and S4). These observations indicate that aerial roots, while necessary, are not sufficient on their own for effective BNF in maize, pointing to the importance of additional factors. Environmental moisture availability, in particular, has emerged as a key regulator for mucilage formation and diazotrophic microbial recruitment. Prior transcriptomic analyses have shown that exposure to external water stimulates gene expression related to polysaccharide biosynthesis in maize aerial root tissues (Pankievicz *et al*., 2022), facilitating polysaccharide secretion from root border cells to form the mucilage gel matrix. This moisture dependence was further supported by our greenhouse experiments, where we compared maize grown under aerial misting to those maintained solely with drip irrigation. In mist-treated conditions, mucilage was abundantly produced on aerial roots, whereas it was absent in drip-irrigated controls (Figure 2). These results reinforce the conclusion that high ambient humidity is essential for mucilage secretion. The moisture-dependent nature of mucilage secretion also suggests that prior field studies conducted under dry or inconsistent irrigation may have underestimated both the prevalence and genetic control of this trait.

While mist irrigation in greenhouse trials reliably stimulated mucilage production, achieving uniformly high humidity in open-field conditions of *Zea* Trial 1 proved challenging. Moreover, the native microbial communities in California’s Central Valley differ substantially from those of the highland environments of Totontepec maize, potentially limiting colonization by compatible diazotrophs. These combined factors may explain the high phenotypic variability observed for NDFA in this study. Based on the moderate heritability for NDFA observed in *Zea* Trial 1, we anticipated detecting genomic regions associated with this trait in the F2:F3 field mapping experiment of *Zea* trial 2. However, NDFA expression proved highly variable across genotypes and no NDFA-associated QTL were detected. Consequently, our genetic mapping analyses focused on aerial roots and other yield component traits, which displayed higher heritability and more consistent expression across the environment, enabling more robust QTL detection.

### QTL Mapping of Aerial Roots and Other Yield Component Traits

Moderate to high heritability estimates for AR and other yield components observed in *Zea* Trial 1 provided a strong rationale to investigate the genetic control for these traits via QTL mapping in the F2:F3 populations of *Zea* Trial 2. The paucity of large effect QTL identified is consistent with the hypothesis that these traits have polygenic architectures, similar to other pre-domestication traits (Buckler *et al*., 2009). The absence of coincident QTL across the three bi-parental subpopulations for any given trait may reflect the relatively small sizes of each subpopulation and corresponding limited statistical power to detect QTL of minor effect. In addition, the lower mapping resolution for certain QTL is likely a consequence of the small population sizes and the limited number of informative recombinant genotypes, which can lead to broad confidence intervals for QTL positions. Future genetic mapping efforts with larger populations, higher marker density, and possibly multi-environment trials will improve the power to detect small-effect QTL and increase mapping resolution.

### QTL Comparison to Published Literature

This study identified both novel and previously reported QTL for AR and other yield-related traits. Overlap between QTL detected here and those described in prior studies provides an important form of validation, while novel QTL expand the existing knowledge of genetic architecture underlying these traits. Each QTL reported in this work is listed by its designated ‘QTL Name’ in Table 2, along with positional information and overlap with previous reports where applicable.

Focusing first on aerial root nodes, AR QTL 1.4 (293.3 – 295.3 Mb) identified in this study is proximal to, but does not overlap with, QTL reported by Zhang *et al*. (2018) for ‘root number’ (286 – 290.8 Mb), ‘aerial root number’ (283.2 – 290.3 Mb), and ‘root layer number’ (285.1 – 290.2 Mb). AR QTL 9.3 (131.6 – 143.8 Mb) overlaps with the previously reported QTL for ‘crown root layer number’ (134.8 – 149.2 Mb) and is proximal to another QTL for ‘aerial root layer number’ (144.8 – 152.5 Mb) (Zhang *et al*., 2018). AR QTL 9.2 (33.7 – 111.4 Mb) appears to be novel, with no published reports of QTL in this defined region associated with aerial root trait.

In addition, we identified a flowering time QTL DPS QTL 10.1 (100.3 – 129.0 Mb), that coincides with the *ZmCCT* locus, a well-characterized regulator of photoperiod sensitivity and flowering time in maize (Hung *et al*., 2012). Notably, Zhang *et al*. (2018) also reported that the *ZmCCT* region is coincident with QTL for ‘crown root layer number’, ‘crown root number’, and ‘crown root number per layer’, suggesting potential pleiotropic effects or close linkage between flowering time regulation and root development. Such co-localization of developmental and architectural traits has been observed in other cereals, where flowering time loci can influence vegetative growth duration and resource allocation to root systems (Wingler *et al*., 2025).

Closer inspection of AR QTL 1.4 using MaizeGDB (Portwood *et al*., 2019) reveals that this relatively narrow interval contains 178 annotated gene models, including confirmed protein coding genes, transposable elements, and low-confidence genes. According to Table S4, ten confirmed protein-coding genes in this QTL exhibit upregulated protein expression in tissues relevant to root development in young seedlings, including root cortex, elongation zone, meristem zone, primary roots, secondary roots, and/or the root stele. Of particular interest are the two uncharacterized protein coding genes, GRMZM2G055834 and GRMZM2G081105, which show RNA transcript upregulation in crown roots at nodes 1-4 and nodes 1-6, respectively (Stelpflug *et al*., 2016; Portwood *et al*., 2019). Future work should prioritize fine-mapping of this region to narrow down candidate genes and explore their functional roles in aerial root development. Targeted mutagenesis approaches, such as CRISPR-Cas9 knockouts or promoter editing, combined with high-resolution phenotyping for aerial root number, node position, and mucilage production, could help clarify the biological roles of these candidate loci.

Finally, during the writing of this work, Laspisa *et al*. (2025) published a similar analysis of aerial root QTL using different germplasm. Some of their QTL may overlap with those published here (particularly AR QTL 9.2 and 9.3), though another of our QTL (AR QTL 1.4) appears distinct. Analysis of the overlap of these QTL to narrow down intervals, and evaluation of *mexicana* admixture in additional material, would be productive avenues of future research.

### Admixture Analysis of *mexicana* into Totontepec Maize

Our phenotypic analyses revealed that *mexicana* produces larger aerial roots than both *parviglumis* and *diploperennis*, as well as more above-ground root-bearing nodes than *parviglumis*. Morphologically, the aerial root phenotype of *mexicana* is more similar to that of Totontepec maize than other wild *Zea* taxa. Based on these observations, we hypothesized that the capacity of abundant aerial root formation and mucilage secretion in Totontepec maize and other traditional varieties may be the result of historical introgression from *mexicana*.

To test this hypothesis, we quantified the extent of *mexicana* introgression in Totontepec maize and examined the overlap between high-introgression genomic regions and the QTL identified in this study. Previous population genomic studies have documented substantial admixture between highland maize and *mexicana* in multiple regions of Mexico (Ross-Ibarra *et al*., 2009; van Heerwaarden *et al*., 2010; Hufford *et al*., 2013), with evidence that such introgression can contribute to local adaptation, particularly to highland environments characterized by cooler temperatures, reduced oxygen partial pressure, and variable moisture regimes (Calfee *et al*., 2021; Barnes *et al*., 2022). Notably, *mexicana* itself displays phenotypic variation for traits beneficial in these environments, including pronounced aerial root development and high mucilage production capacity, traits that may facilitate BNF through diazotroph recruitment.

Our admixture analysis revealed that *mexicana*-derived introgression segments in Totontepec maize are not randomly associated with the AR and SC QTL identified here, with a greater degree of overlap than expected under a uniform genomic distribution (Figure 4, Table S2). These results highlight multiple candidate genomic regions where *mexicana*-derived alleles may contribute to yield-related and structural traits in Totontepec maize. We note that while there also was overlap between *mexicana*-derived introgression segments and QTL for DPS and PDM, this was not greater than would be expected by chance.

The striking morphological similarity in aerial root structure and mucilage secretion capacity between Totontepec maize and *mexicana*, combined with the non-random overlap of introgression segments and relevant QTL, provide compelling evidence for a potential case of adaptive introgression. Similar cases have been reported in which introgression from wild relatives has contributed to beneficial agronomic traits in crops (Janzen *et al*., 2019), supporting the possibility that *mexicana*-derived alleles play a role in preserving or strengthening traits associated with BNF in Totontepec maize. This hypothesis should be tested in follow-up investigations, potentially including fine-mapping of the traits using larger mapping populations, as well as comparative genomic analyses between *mexicana* and Totontepec maize.

### Conclusions and Future Directions

Previous studies have demonstrated that the aerial roots and associated mucilage of Totontepec maize can facilitate the recruitment of diazotrophic bacteria, contributing to BNF. In this work, we quantified the heritability of AR traits, NDFA, and other yield-related traits, and identified QTL associated with AR and multiple agronomic traits. We also present preliminary evidence of overlap between these QTL and genomic regions with high levels of *mexicana* introgression, raising the possibility that mexicana-derived alleles contribute to or even have originated BNF-related traits in Totontepec maize.

Identified QTL, particularly those overlapping with introgressed regions, require further fine-mapping to narrow candidate intervals and pinpoint to causal genes. Precision phenotyping of mapping populations under controlled, uniformly moist conditions that mimic the native highland environment may strengthen the ability to detect genetic loci influencing aerial root abundance, mucilage production, and NDFA. For the relatively narrow AR QTL 1.4, which contains 11 candidate protein-coding genes, we recommend functional validation using targeted approaches such as CRISPR-Cas9 mutagenesis or transgenic overexpression.

## DATA AVAILABILITY

Main and supplemental figures and tables have been deposited via the GSA figshare portal under the file names ‘2025_Totontepec_QTL_Manuscript_Figures-main_Final’ and ‘2025_Totontepec_QTL_Manuscript_Figures-supplementary_Final’; both files were submitted under the title ‘2025 – QTL for AR and Yield Traits in Zea mays – Manuscript Figures – ODonnell-et-al’. Datasets and code used for analyses have been uploaded to the corresponding Github repository <https://github.com/daodonnell12/qtl-aerial-roots-yield-traits-zea-mays>. Genetic resources, including biological materials and nucleic acid sequences, were accessed under Access and Benefit Sharing (ABS) Agreements between the Community of Totontepec Villa De Morelos, Mixe, Oaxaca and Mars, Incorporated, and with authorization from the Mexican government. An internationally recognized certificate of compliance has been issued by the Mexican government under the Nagoya Protocol for such activities (ABSCH-IRCC-MX-207343-3). Any party seeking access to the genetic resources underlying the analysis reported here should obtain the authorization from the government of Mexico.

## ACKNOWLEDGEMENTS

The authors of this publication thank the community of Totontepec Villa de Morelos for their collaboration and support, and particularly Maria del Refugio Vasquez Cano for assistance with seed collections. We also thank the UC Davis Plant Sciences Core Field and Greenhouse teams for support of all trials presented in this work, and Cornell University for Genotyping-by-Sequencing services. This research used the High-Performance Computing Core Facility (HPC@UCD) at the University of California, Davis.

## STUDY FUNDING

The research was supported by grants from BioN2 and by the Foundation for Food and Agriculture Research grant #CA19-SS-0000000100. JR-I would like to acknowledge support from the US Dept. of Agriculture (Hatch project CA-D-PLS-2066-H 548).

## Notes

https://github.com/daodonnell12/qtl-aerial-roots-yield-traits-zea-mays

### Competing Interest Statement

The authors have declared no competing interest.

### Summary of Updates

The original manuscript submission did not contain the main figures or supplementary figures/tables due to a file upload error stemming from oversized files. The new submission includes the main and supplementary figures.

