## Supplementary Material for "HERITABILITY AND QTL MAPPING OF AERIAL ROOTS AND OTHER YIELD COMPONENT TRAITS WITH IMPLICATIONS FOR N_2_ FIXATION IN *ZEA MAYS*"

**Supplementary Figure 1:** Map of Experimental Entries in *Zea* Trial 1

**a**

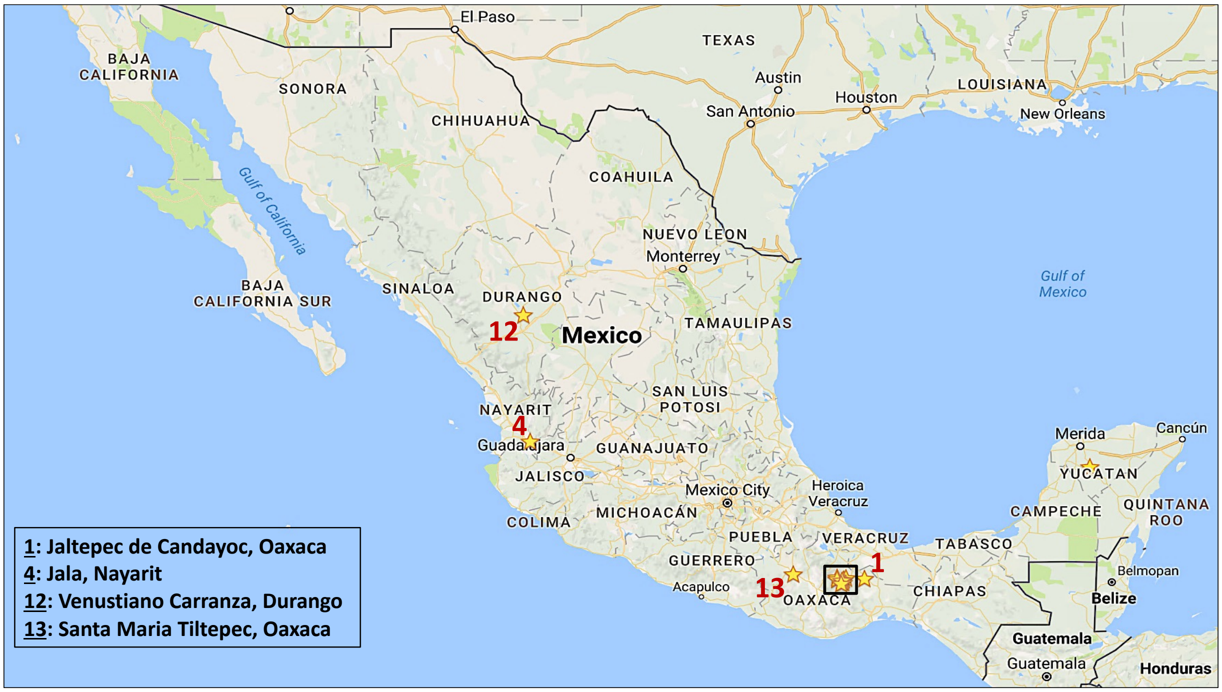


**b**

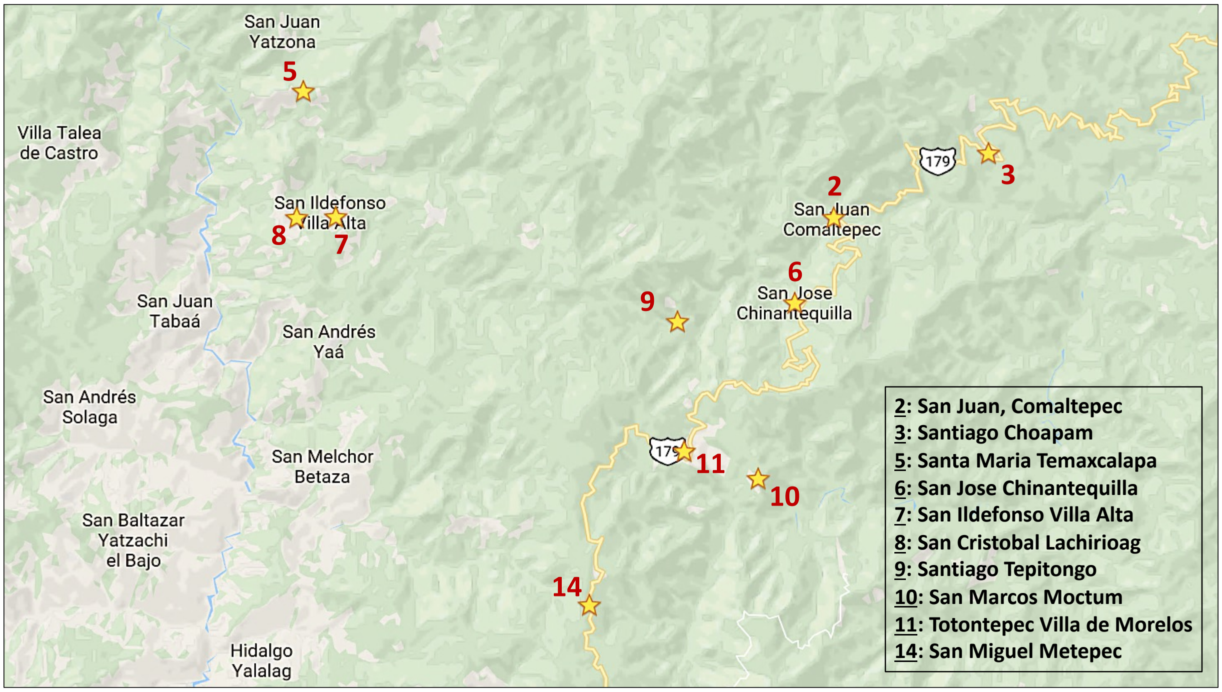


**Supplementary Figure 1:** Map panels showing experimental entries of *Zea* Trial 1, corresponding to the list order indicated in Table S1 (column 2 'Map Position'). Panel '**a**' shows a full view of entries throughout Mexico, while panel '**b**' zooms in on panel A's box outline of closely clustered entries within Oaxaca. Panels obtained from Google Maps (Google).

**Supplementary Figure 2:** *Zea* Trial 1 Soil Analyses – Nutrient Profile and $^{15}N$

**a**

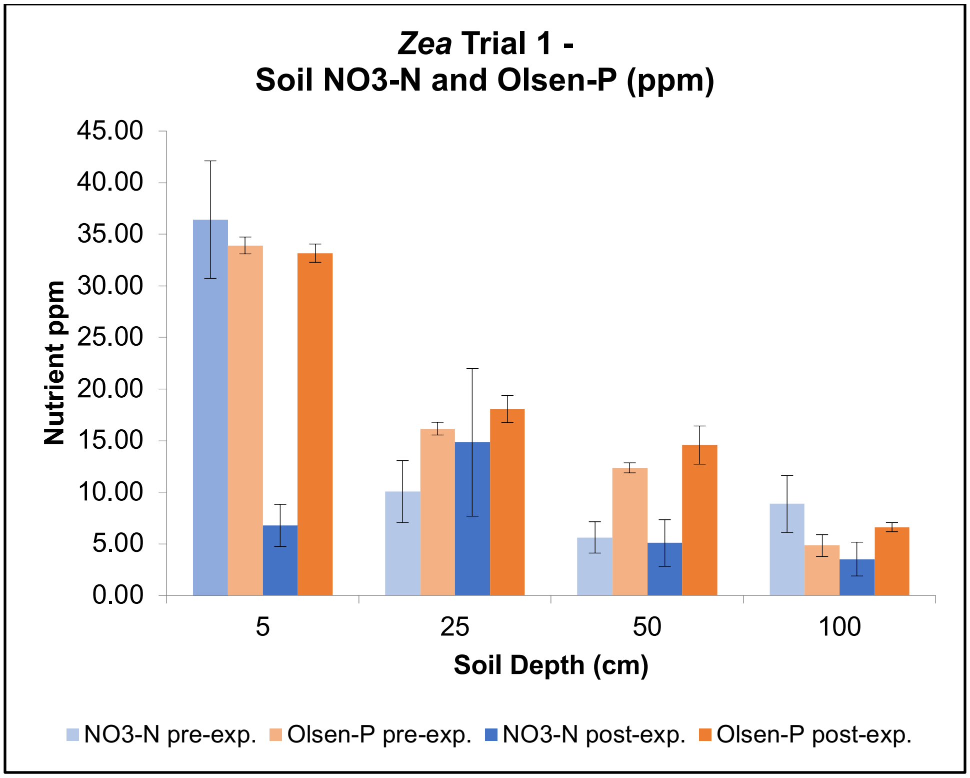


**b**

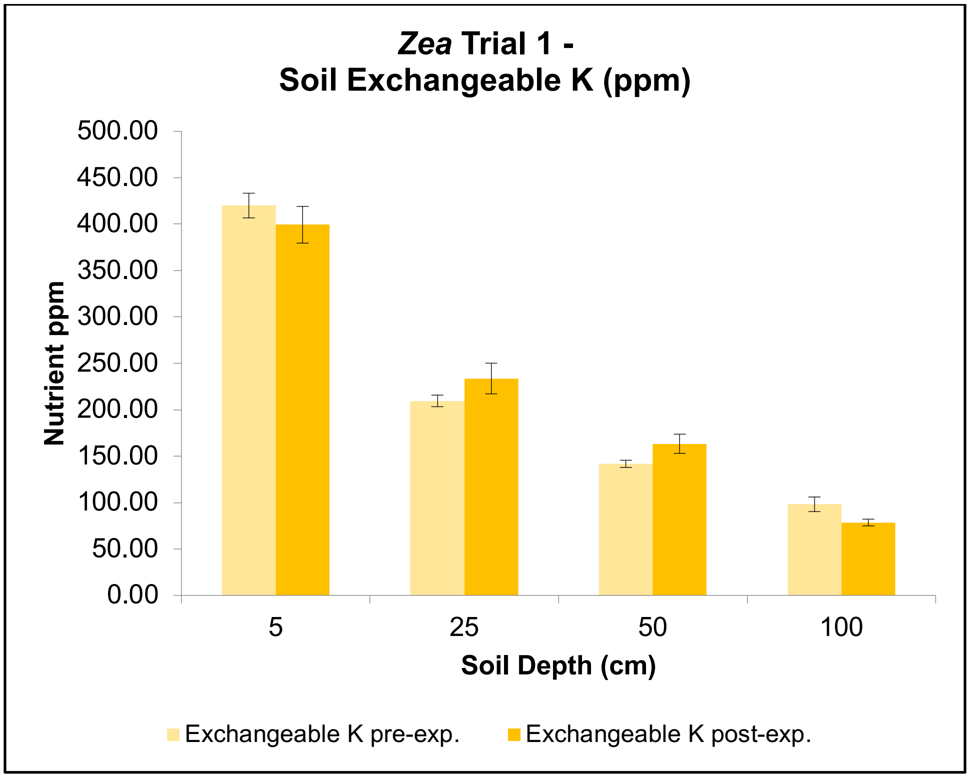

**c**

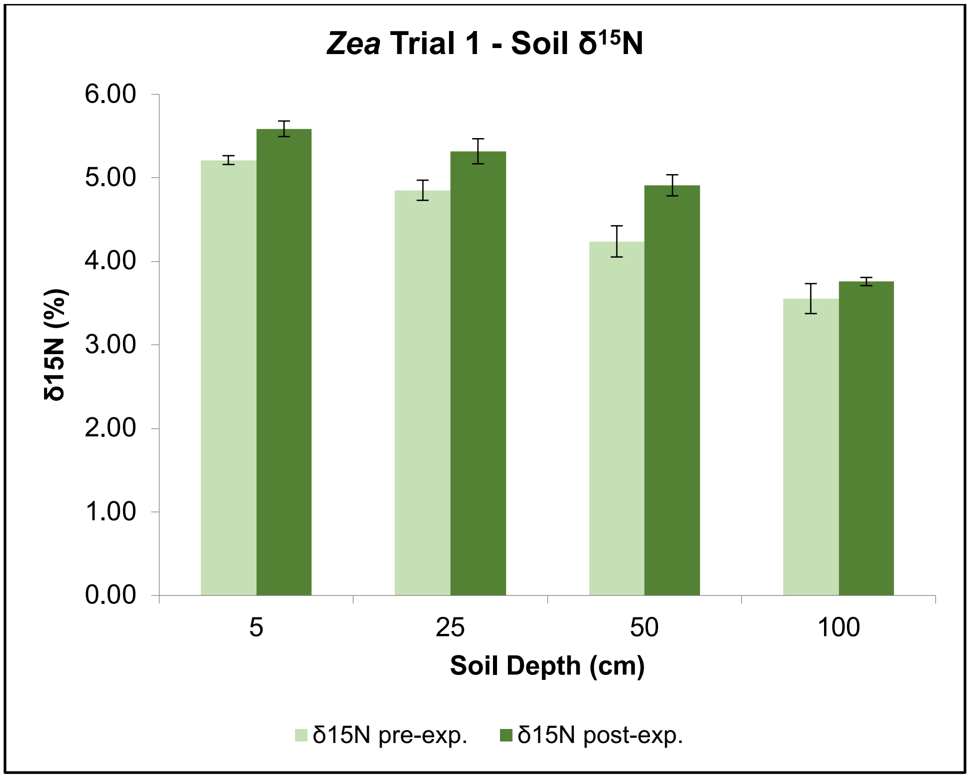


**Supplementary Figure 2:** For *Zea* Trial 1, soil analyses for the field in which plants were grown. From top to bottom: (**a**) $NO_3$-N and Olsen-P, (**b**) Exchangeable K, and (**c**) $^{15}N$ soil nutrient analysis for soil sampled prior to and after experiment. For fifteen sampled locations evenly distributed throughout the field pre-experiment and four sampled locations post-experiment, values were averaged per sampling depth (5, 25, 50, and 100 cm.). X-axis = Soil Depth (cm.). Y-Axis = Nutrient ppm value or $\delta^{15}N$ % value. Legend color keys are indicated at bottom of each chart, with abbreviations 'pre-exp.' and 'post-exp.' to indicate values measured before and after experiment, respectively. Standard error bars are shown.

**Supplementary Figure 3:** NDFA Variation among *Zea* Subpopulations

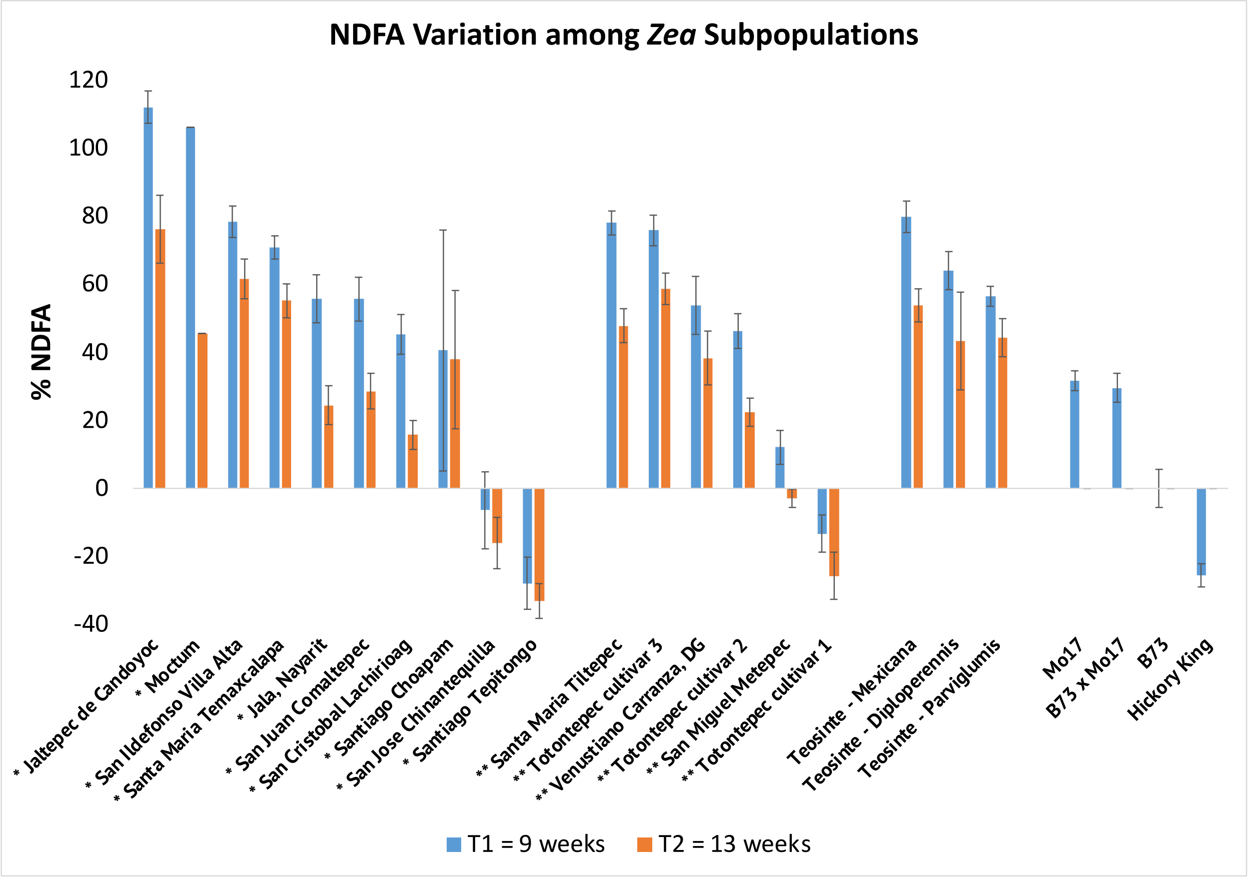


**Supplementary Figure 3:** %NDFA values corresponding to *Zea* Trial 1 across two time-points, calculated using B73 improved cultivar as the reference negative control. Entries are indicated along the X-axis and organized first by *Zea* group (left to right: lowland maize, highland maize, teosinte, improved cultivar) and then by %NDFA value from greatest to least within each group. %NDFA values are indicated via the Y-axis. Color Key: blue bars = time-point 1 (9 weeks after transplant); orange bars = time-point 2 (13 weeks after transplant). Standard error bars are shown.

**Supplementary Figure 4:** Aerial Root Node, Height, and Maturity Time Variation among *Zea* Subpopulations

**a**

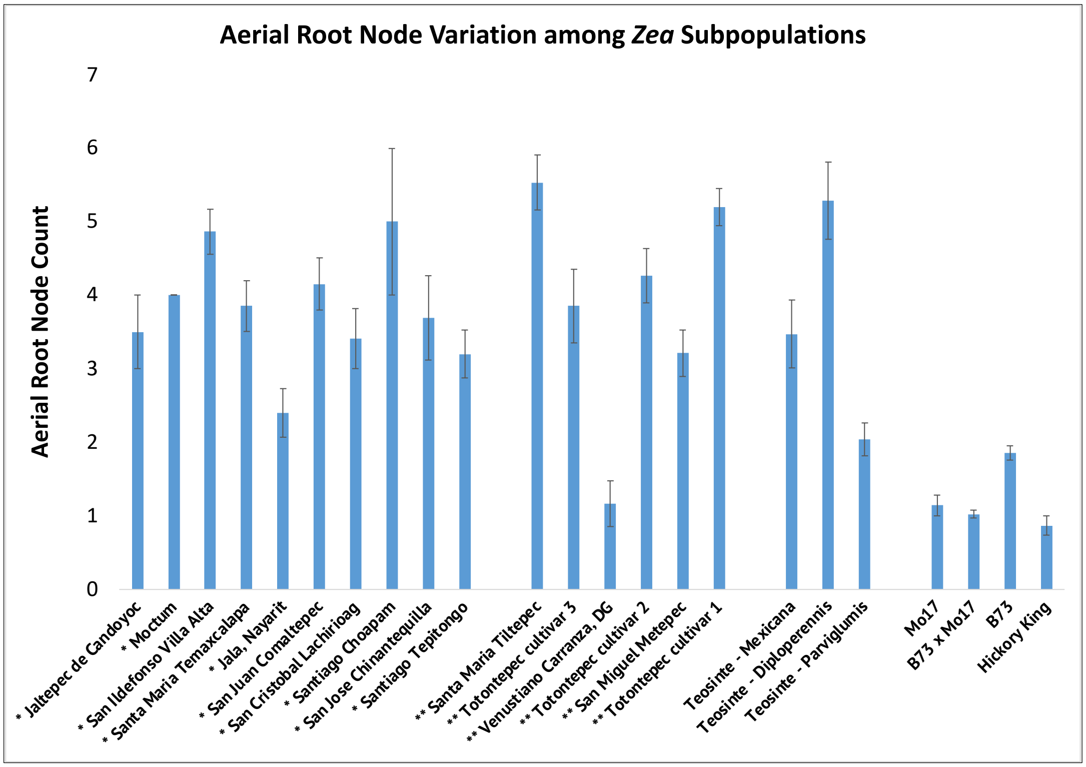


**b**

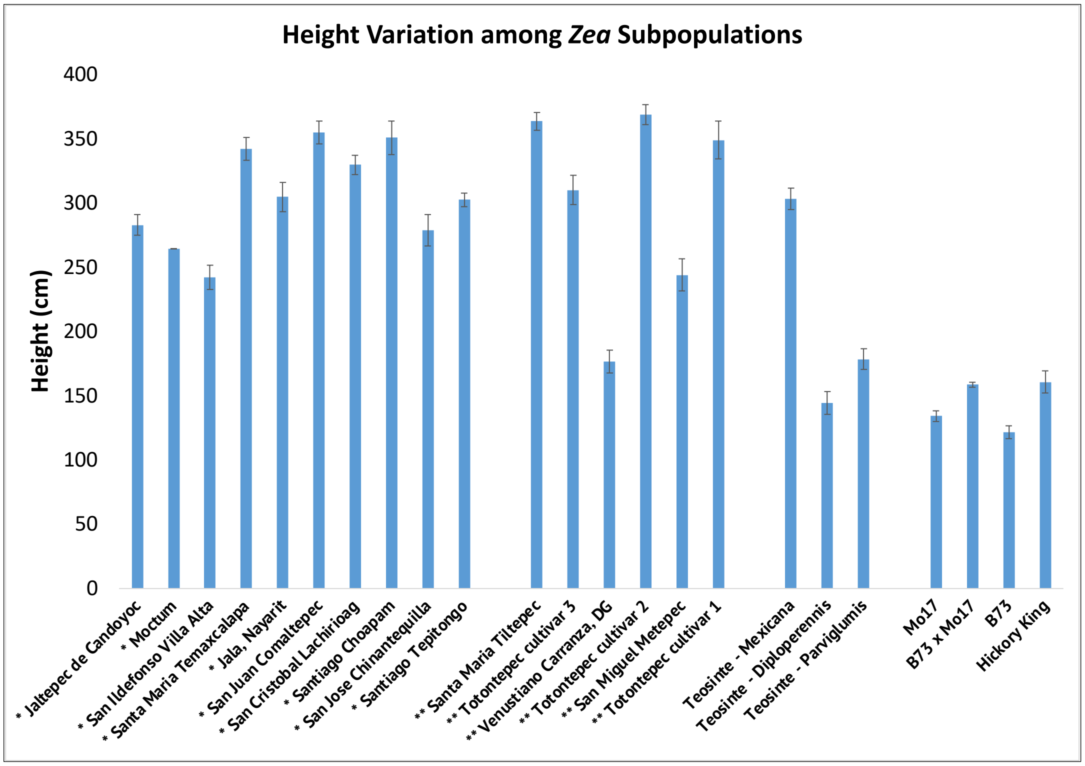

c

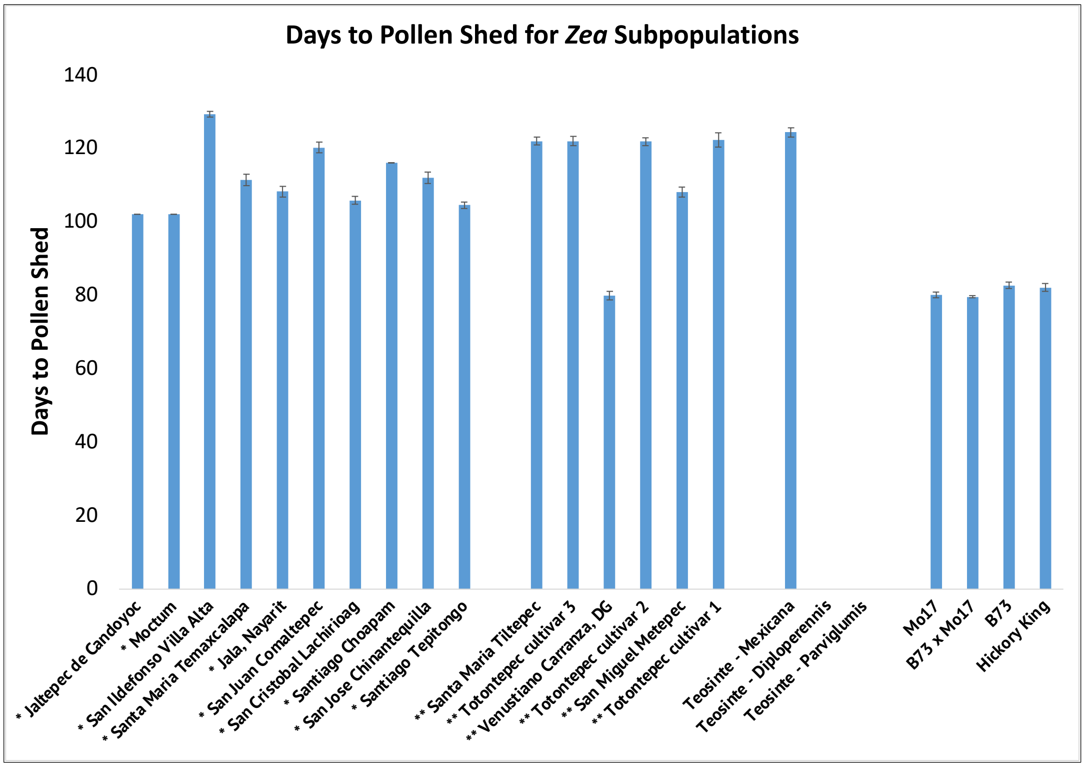


**Supplementary Figure 4:** Bar charts displaying the variation in phenotypes among *Zea* groups previously specified. Entries are displayed in the same order corresponding to Figure S3A. X-axis = entry. Y-axis = corresponding phenotype. From top to bottom: (**a**) number of nodes possessing aerial roots in fully mature plants, (**b**) height (cm.), and (**c**) days to maturity (as assessed via days to pollen shed). Within chart C, *diploperennis* and *parviglumis* do not have values as they did not shed pollen. Standard error bars are shown.

**Supplementary Figure 5:** *Zea* Trial 2 Soil Analyses – Nutrient Profile and $^{15}N$

**a**

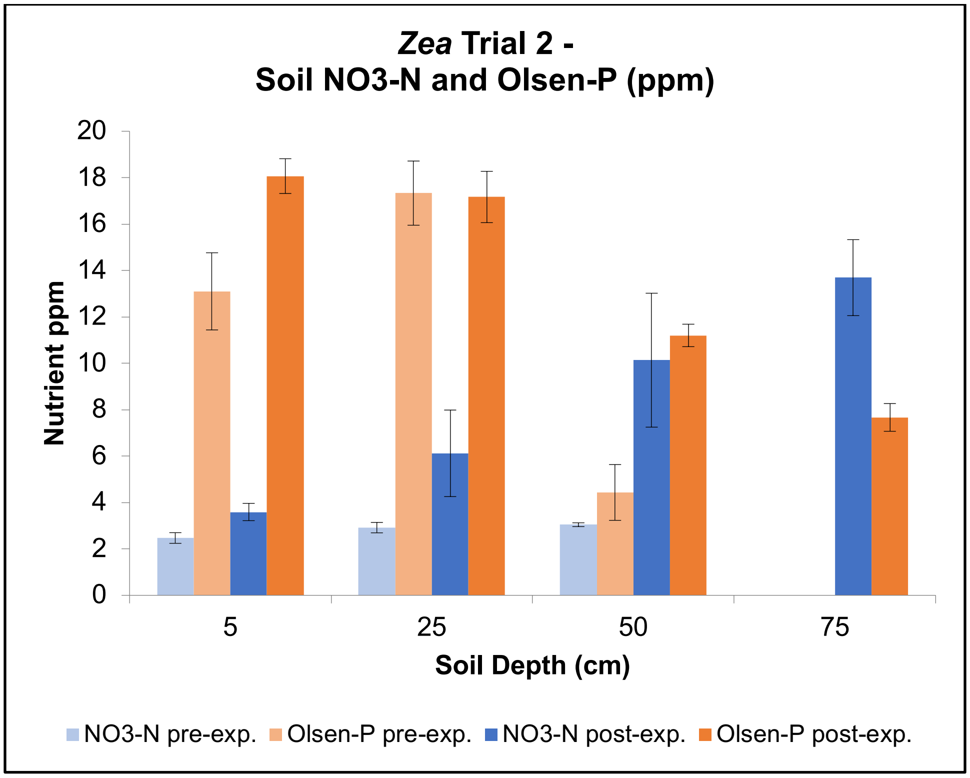


**b**

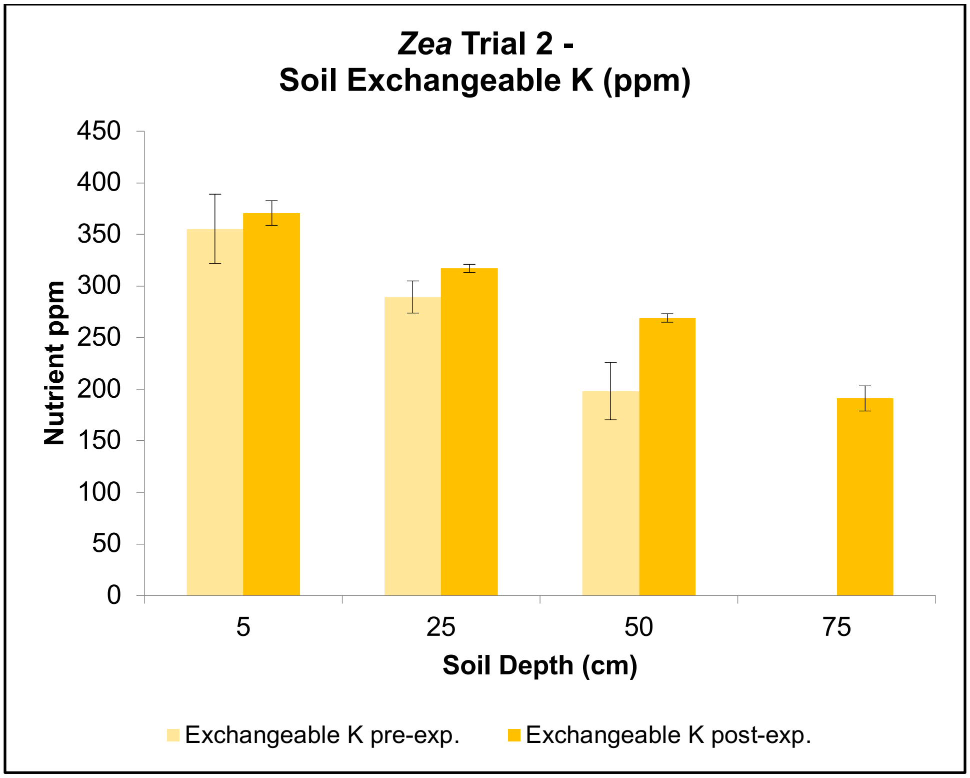

c

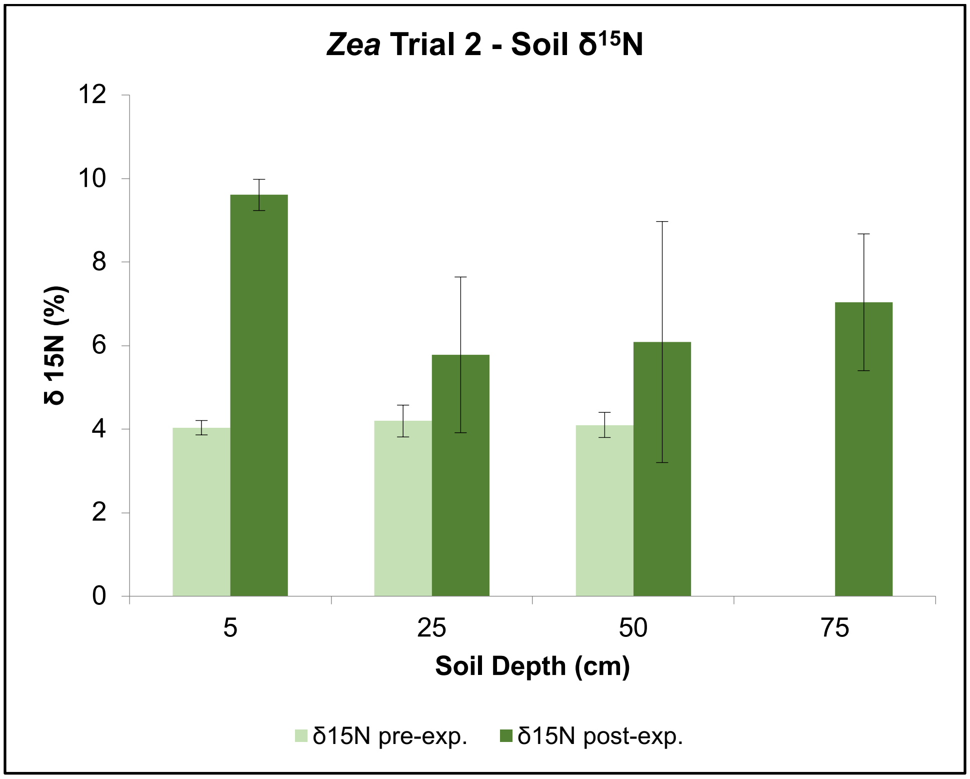


**Supplementary Figure 5:** For *Zea* Trial 2, soil analyses for the field in which plants were grown. From top to bottom: (**a**) $NO_3$-N and Olsen-P, (**b**) Exchangeable K, and (**c**) $^{15}N$ soil nutrient analysis for soil sampled prior to and after experiment. For three sampled locations evenly distributed throughout the field pre-experiment and nine sampled locations post-experiment, values were averaged per sampling depth (5, 25, 50, and 100 cm.). X-axis = Soil Depth (cm.). Y-Axis = Nutrient ppm value or $\delta^{15}N$ % value. Legend color keys are indicated at bottom of each chart, with abbreviations 'pre-exp.' and 'post-exp.' to indicate values measured before and after experiment, respectively. Standard error bars are shown.

**Supplementary Figure 6:** *Zea* Trial 2 Phenotype Distributions for Traits of Interest

**a**

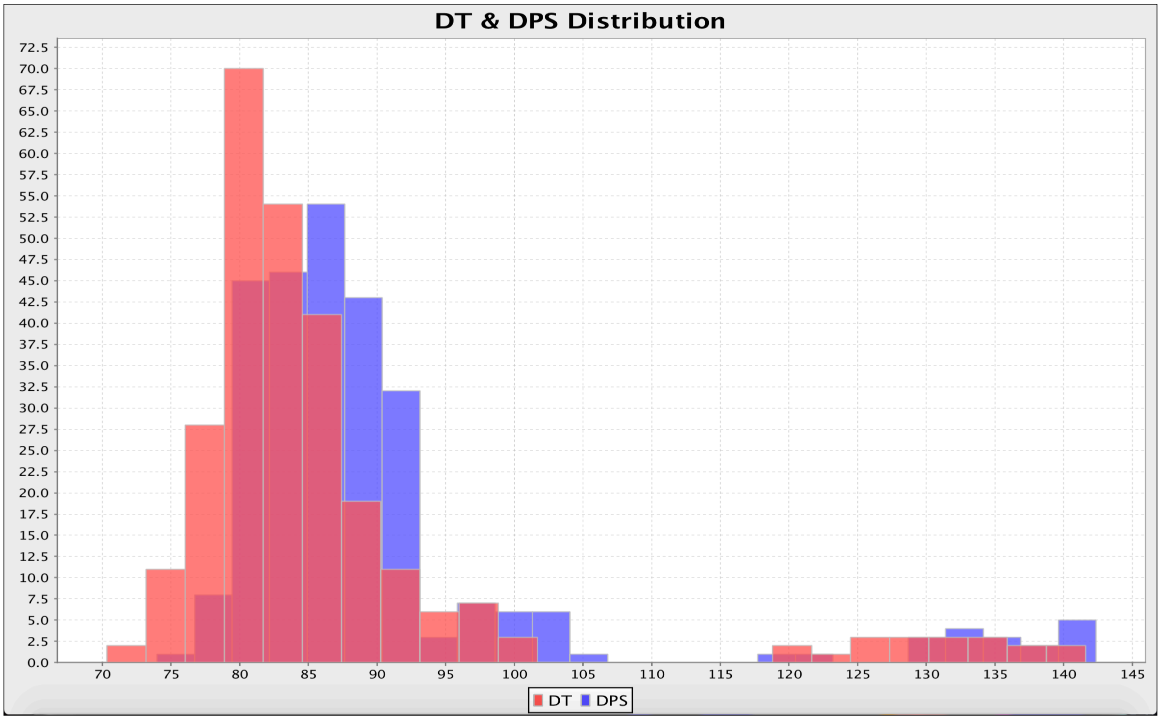


**b**

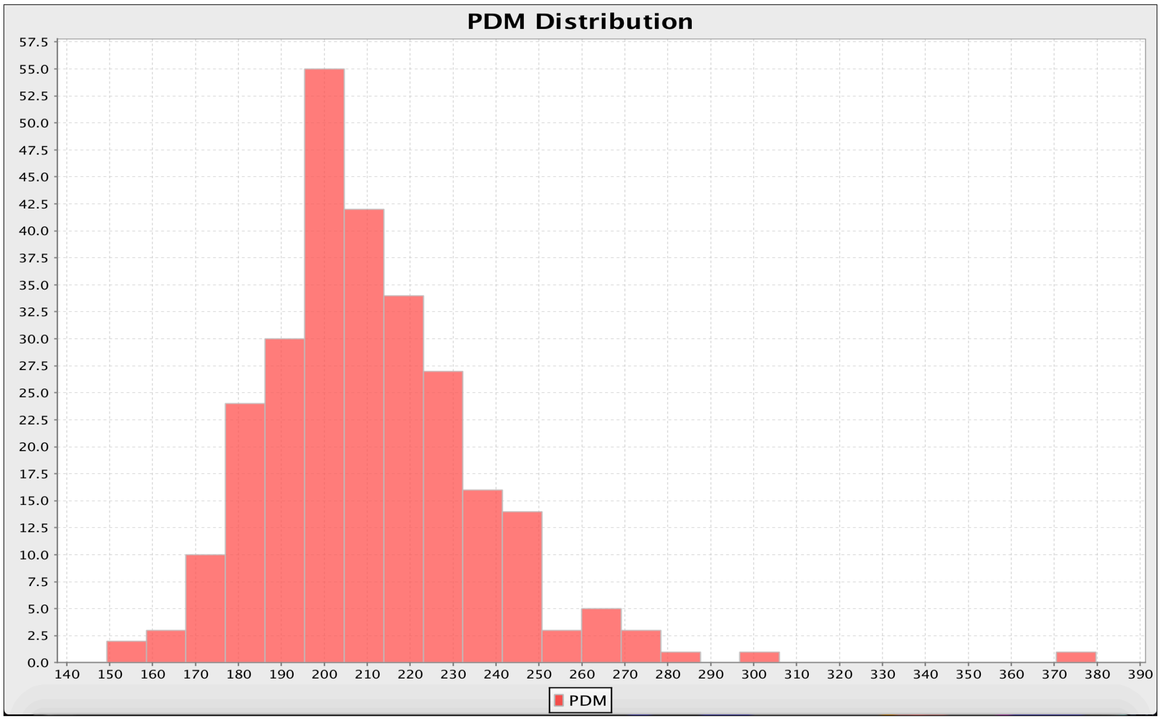

c

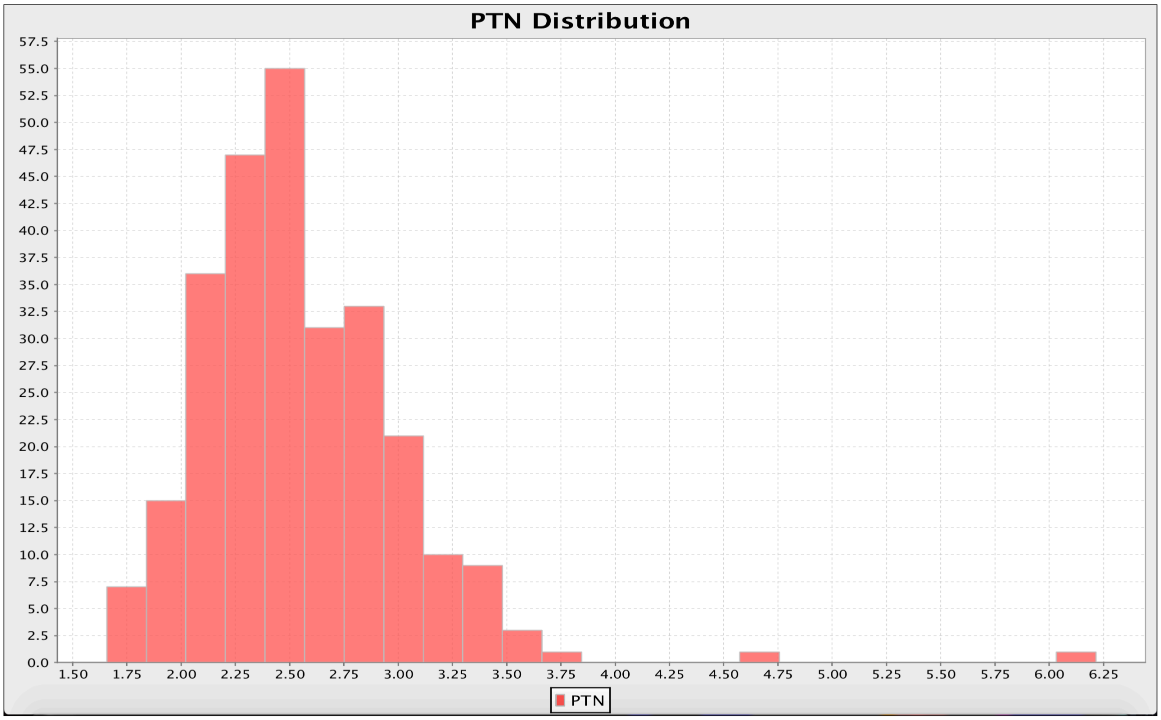


d

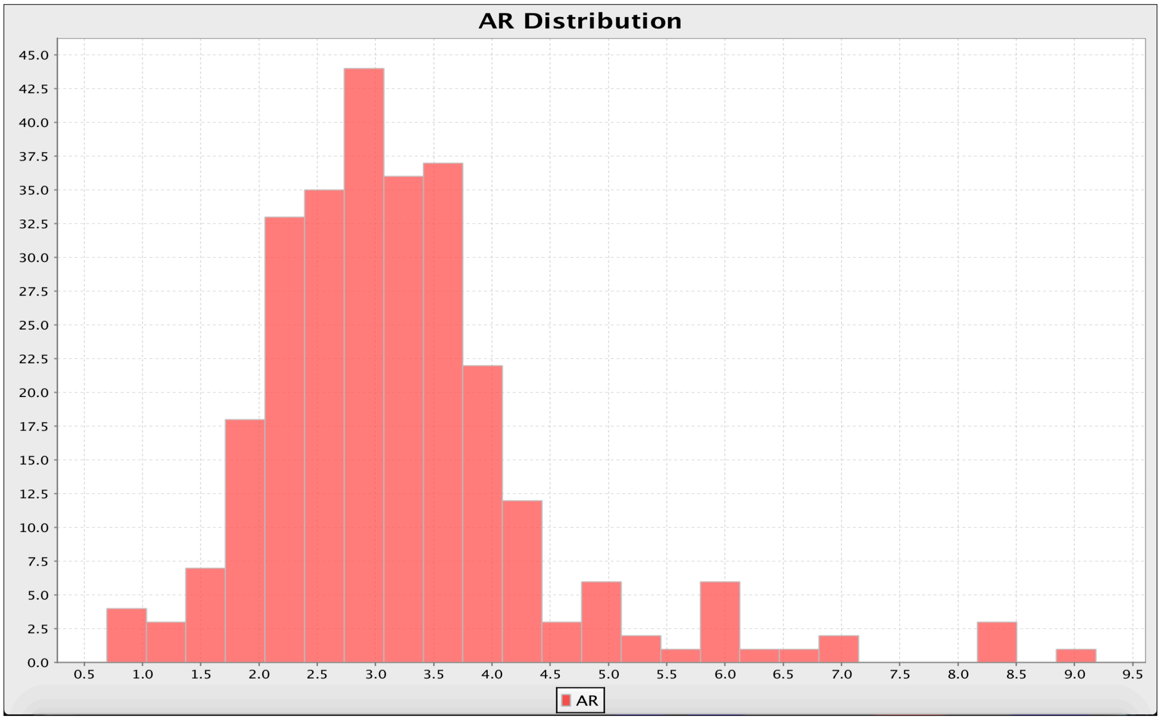


**Supplementary Figure 6:** *Zea* Trial 2 trait histograms, from top to bottom, for (**a**) time to maturity [DT = days to tassel emergence; DPS = days to pollen shed], (**b**) mature plant dry mass [PDM], (**c**) mature plant total nitrogen [PTN], and (**d**) aerial root node count [AR]. X-axis = trait value bin. Y-axis = frequency of trait value within population.

**Supplementary Figure 7:** Genetic Maps per Bi-parental Population of *Zea* Trial 2

**a**

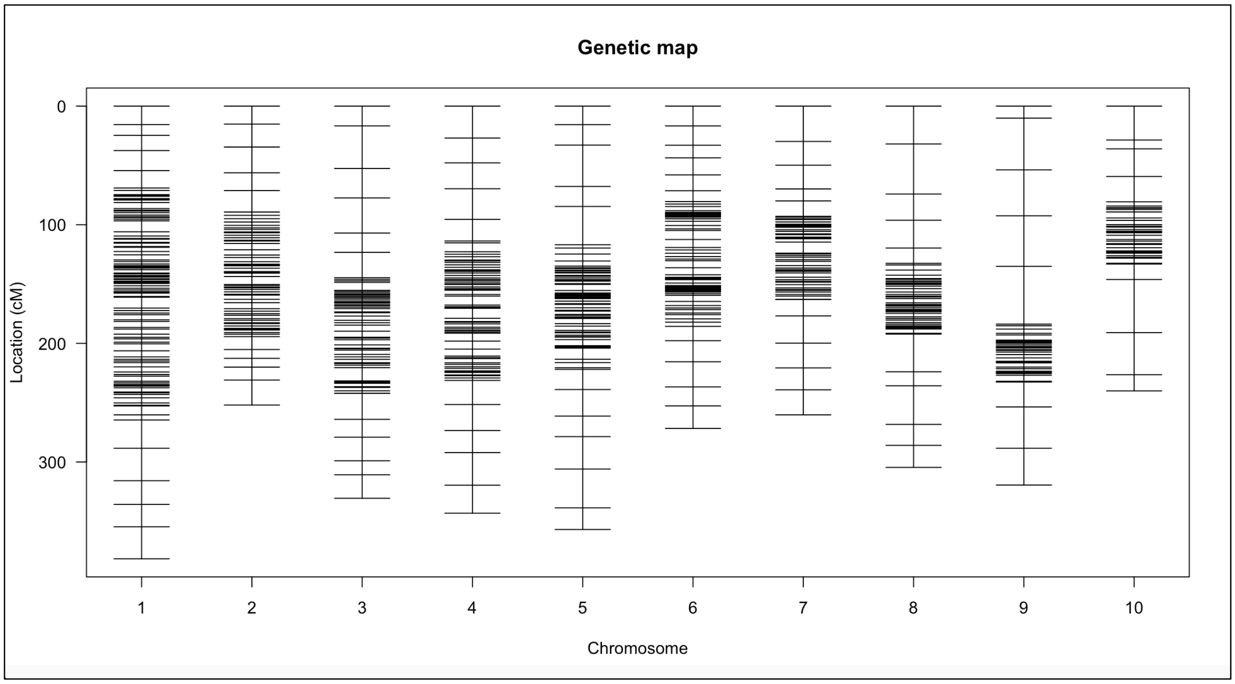


**b**

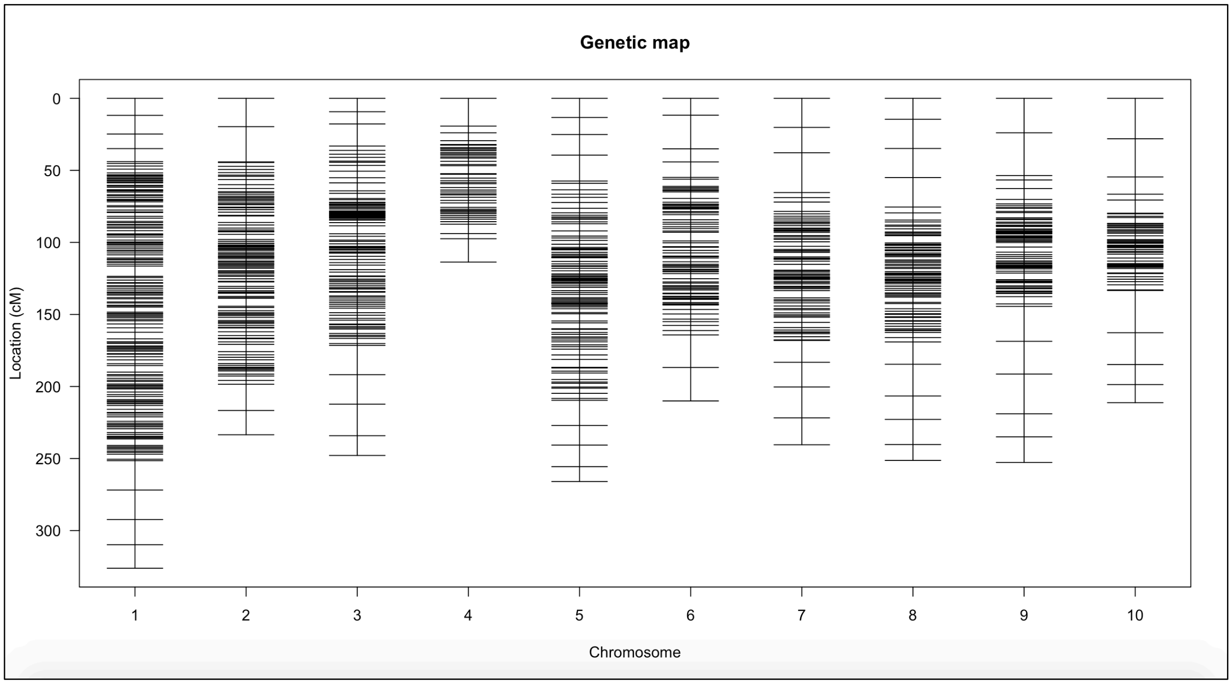

**c**

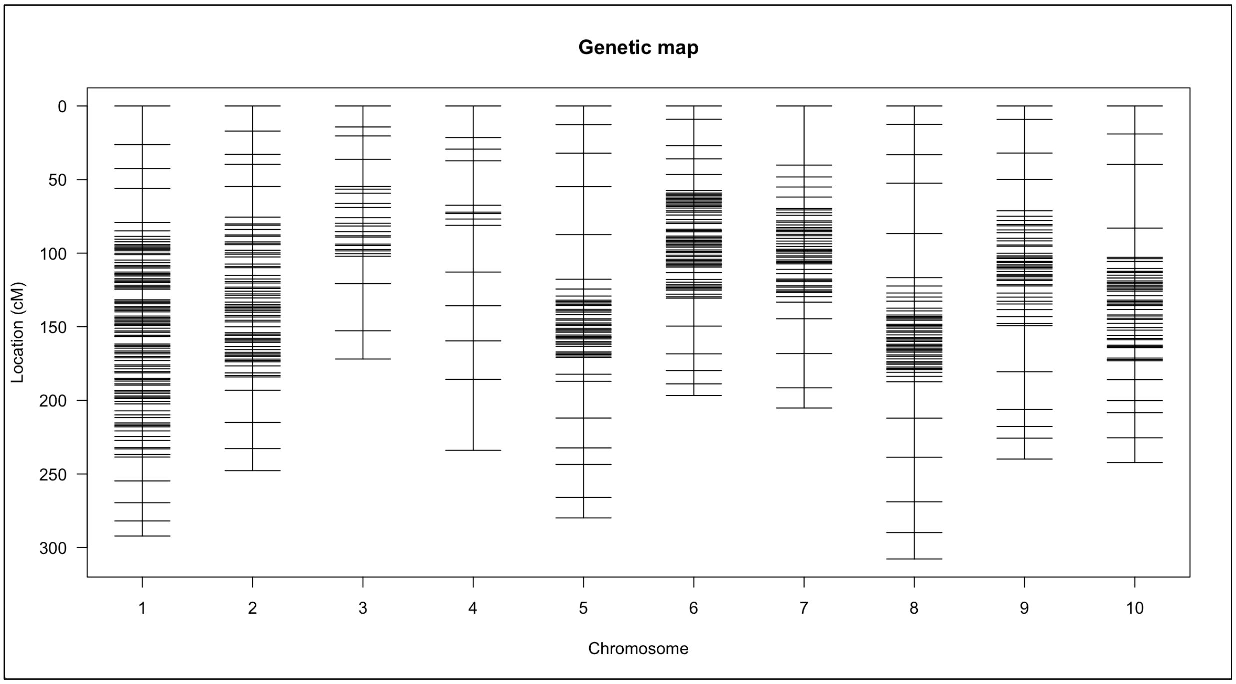


**Supplementary Figure 7:** Genetic maps per bi-parental subgroup of *Zea* Trial 2. From top to bottom: (**a**) Totontepec x B73, (**b**) Totontepec x LH82, and (**c**) Totontepec x Mo17. X-axis = chromosome number. Y-axis = location of markers (centimorgans distance).

**<u>Supplementary Table 1:</u>** <u>Entries of *Zea* Trial 1</u>

**Supplementary Table 1:** Columns from left to right: (1) All entries of *Zea* Trial 1, including traditional cultivars indicated by location of origin with Mexican province abbreviation, if applicable; (2) Map Position corresponding to S. Figure S1; (3) USDA accession ID, if applicable ('NA' entries were collected in-situ); (4) evolutionary status, color-coded, wherein pre-domesticate = purple, post-domesticate lowland traditional cultivar = green, post-domesticate highland traditional cultivar = red, and improved types = blue); (5) natural elevation; and (6) sample size for the trial. *Z. diploperennis* is listed with an asterisk (*) to designate as a separate species of genus *Zea*. Lowland cultivars are considered as those of regions <1,600 m elevation, while highland cultivars are those of regions >1,600 m elevation. Inbred line elevation not applicable due to recent development within the past century. Elevation information for traditional cultivars was obtained from <mexico.pueblosamerica.com> (Mexico.PueblosAmerica.com, 2023). [Accessed November 09, 2023].

| ***Zea* Entry** | **Map Position (S. Figure S1)** | **Accession ID** | **Evolutionary Status** | **Origin Elevation** | **Sample Size** |
|---|---|---|---|---|---|
| *Z. diploperennis** | NA | PI 462368 | Pre-domestication | 1,400 – 2,400 m | 8 |
| *Z. mays* spp. *parviglumis* | NA | Ames 21809 | Pre-domestication | < 1,800 m | 34 |
| *Z. mays* spp. *mexicana* | NA | PI 566674 | Pre-domestication | > 1,700 m | 27 |
| Jaltepec de Candoyoc, OA | 1 | NA | Post-domestication | 62 m | 3 |
| San Juan Comaltepec, OA | 2 | NA | Post-domestication | 609 m | 24 |
| Santiago Choapam, OA | 3 | NA | Post-domestication | 879 m | 3 |
| Jala, NA | 4 | PI 645922 | Post-domestication | 1,065 m | 25 |
| Santa Maria Temaxcalapa, OA | 5 | NA | Post-domestication | 1,082 m | 28 |
| San Jose Chinantequilla, OA | 6 | NA | Post-domestication | 1,171 m | 25 |
| San Ildefonso Villa Alta, OA | 7 | NA | Post-domestication | 1,202 m | 19 |
| San Cristobal Lachirioag, OA | 8 | NA | Post-domestication | 1,227 m | 25 |
| Santiago Tepitongo, OA | 9 | NA | Post-domestication | 1,542 m | 25 |
| San Marcos Moctum, OA | 10 | NA | Post-domestication | 1,558 m | 1 |
| Totontepec cultivar 1, OA | 11 | NA | Post-domestication | 1,821 m | 14 |
| Totontepec cultivar 2, OA | 11 | NA | Post-domestication | 1,821 m | 26 |
| Totontepec cultivar 3, OA | 11 | NA | Post-domestication | 1,821 m | 21 |
| Venustiano Carranza, DG | 12 | NA | Post-domestication | 1,968 m | 6 |
| Santa Maria Tiltepec, OA | 13 | NA | Post-domestication | 2,179 m | 24 |
| San Miguel Metepec, OA | 14 | NA | Post-domestication | 2,641 m | 24 |
| B73 inbred line | NA | PI 550473 | Improved line | N/A | 27 |
| Mo17 inbred line | NA | PI 558532 | Improved line | N/A | 22 |
| Hickory King | NA | PI 482981 | Improved line | N/A | 28 |
| B73 x Mo17 hybrid | NA | Ames 19097 | Improved hybrid | N/A | 155 |

**Supplementary Table 2:** QTL Enriched w/ Elevated *Mexicana* Introgression into Totontepec Maize

**Supplementary Table 2:** This table shows only QTL regions enriched with elevated *mexicana* introgression (>0.5) into Totontepec maize. From left to right, columns show: chromosome number, introgression region start point (bp), introgression region end point (bp), proportion of introgression from *mexicana* (0 – 1), total genes in introgression segment, overlapping trait QTL, population in which overlapping trait QTL was identified (wherein 'T' = Totontepec cultivar parent), QTL LOD interval start point (bp), QTL LOD interval end point (bp), total genes in QTL region, percentage of QTL overlapping with the corresponding introgression segment (based on gene overlap). An additional row beneath each unique QTL summarizes the total percentage of the QTL overlapping with elevated introgression segments. Note*: Three introgression regions of chromosome 9 are listed twice (marked with an '*' in column 1), to calculate respective overlap with two coincident trait QTL (AR 9.2 and SC 9.1). Note**: Certain elevated introgression segments are only partially overlapping with the corresponding QTL region. While the total number of genes in the introgression segment are listed in column 5, numbers marked with '**' indicate that only some of those genes fall within the QTL. QTL enrichment (far right column) is calculated based on the number of genes in introgression segments which overlap with the corresponding QTL.

| chr. | intr. region start (bp) | intr. region end (bp) | intr. *mexicana* | total genes in introgression segment | overlapping trait QTL | pop. of identified QTL | LOD interval start (bp) | LOD interval end (bp) | total genes in QTL | % of QTL enriched w/ introgression |
|---|---|---|---|---|---|---|---|---|---|---|
| 3 | 35750177 | 35750177 | 0.502 | 1 | SC (3.1) | T x B73 | 10683505 | 162587460 | 6355 | 0.02% |
| 3 | 38602012 | 41403869 | 0.518 | 124 | SC (3.1) | T x B73 | 10683505 | 162587460 | 6355 | 1.95% |
| 3 | 44095158 | 47574787 | 0.534 | 139 | SC (3.1) | T x B73 | 10683505 | 162587460 | 6355 | 2.19% |
| 3 | 49562562 | 53675536 | 0.623 | 176 | SC (3.1) | T x B73 | 10683505 | 162587460 | 6355 | 2.77% |
| 3 | 63108209 | 66312613 | 0.571 | 101 | SC (3.1) | T x B73 | 10683505 | 162587460 | 6355 | 1.59% |
| 3 | 112098658 | 114790935 | 0.64 | 97 | SC (3.1) | T x B73 | 10683505 | 162587460 | 6355 | 1.53% |
| 3 | 129291770 | 132483085 | 0.594 | 150 | SC (3.1) | T x B73 | 10683505 | 162587460 | 6355 | 2.36% |
| 3 | 136908028 | 139736412 | 0.566 | 137 | SC (3.1) | T x B73 | 10683505 | 162587460 | 6355 | 2.16% |
| 3 | 162297619 | 165022441 | 0.605 | 115** | SC (3.1) | T x B73 | 10683505 | 162587460 | 6355 | 0.17% |
| | | | | 1040** | | | | | 6355 | 14.73% |
| 4 | 67020744 | 71704923 | 0.602 | 176** | PDM (4.1) | T x Mo17 | 67143858 | 237313637 | 8430 | 1.93% |
| 4 | 124954610 | 130571862 | 0.795 | 220 | PDM (4.1) | T x Mo17 | 67143858 | 237313637 | 8430 | 2.61% |
| 4 | 131700301 | 142582826 | 0.655 | 419 | PDM (4.1) | T x Mo17 | 67143858 | 237313637 | 8430 | 4.97% |
| 4 | 168739151 | 168739151 | 0.519 | 0 | PDM (4.1) | T x Mo17 | 67143858 | 237313637 | 8430 | 0.00% |
| 4 | 168753601 | 168976868 | 0.527 | 4 | PDM (4.1) | T x Mo17 | 67143858 | 237313637 | 8430 | 0.05% |
| | | | | 819** | | | | | 8430 | 9.56% |
| 8 | 128973724 | 131113149 | 0.503 | 109 | DPS (8.1) | T x Mo17 | 118971709 | 143492696 | 1462 | 7.46% |
| 8 | 136415654 | 138213242 | 0.512 | 96 | DPS (8.1) | T x Mo17 | 118971709 | 143492696 | 1462 | 6.57% |
| | | | | 205 | | | | | 1462 | 14.02% |

| | | | | | | | | | | |
|---|---|---|---|---|---|---|---|---|---|---|
| 9 | 32946981 | 42289811 | 0.884 | 348** | AR (9.2) | T x LH82 | 33670979 | 111431608 | 3342 | 9.04% |
| 9 | 63841178 | 69060908 | 0.547 | 206 | AR (9.2) | T x LH82 | 33670979 | 111431608 | 3342 | 6.16% |
| 9 | 81425959 | 87873575 | 0.667 | 266 | AR (9.2) | T x LH82 | 33670979 | 111431608 | 3342 | 7.96% |
| | | | | 820** | | | | | 3342 | 23.16% |
| 9* | 32946981 | 42289811 | 0.884 | 348 | SC (9.1) | T x Mo17 | 22782253 | 119839322 | 4402 | 7.91% |
| 9* | 63841178 | 69060908 | 0.547 | 206 | SC (9.1) | T x Mo17 | 22782253 | 119839322 | 4402 | 4.68% |
| 9* | 81425959 | 87873575 | 0.667 | 266 | SC (9.1) | T x Mo17 | 22782253 | 119839322 | 4402 | 6.04% |
| | | | | 820 | | | | | 4402 | 18.63% |
| 10 | 108468473 | 112586972 | 0.844 | 213 | DPS (10.1) | T x B73 | 100252518 | 128967696 | 1601 | 13.30% |
| 10 | 121412033 | 123376961 | 0.648 | 113 | DPS (10.1) | T x B73 | 100252518 | 128967696 | 1601 | 7.06% |
| | | | | 326 | | | | | 1601 | 20.36% |

Supplementary Table 3: All QTL and Proportion of Enrichment w/ Elevated *Mexicana* Introgression into Totontepec Maize

**Supplementary Table 3:** This table shows all QTL regions and corresponding percent enrichment with elevated *mexicana* introgression (>0.5) into Totontepec maize. From left to right, columns show: Trait QTL name, QTL position (expressed as "[chromosome number]:[LOD interval start bp]..[LOD interval end bp]"), total genes in the QTL, total overlapping genes within regions of elevated *mexicana* introgression, percentage of QTL enriched with elevated *mexicana* introgression (based on gene overlap). An additional highlighted row beneath each set of trait QTL summarizes the total percentage of all QTL [for the corresponding trait] enriched with elevated *mexicana* introgression (based on gene overlap).

| Trait QTL name | QTL position | Total genes in QTL | Total genes in elevated intr. regions | % QTL enriched w/ introgression |
|---|---|---|---|---|
| AR 1.4 | Chr1:293329598..295309095 | 180 | 0 | 0.00% |
| AR 9.2 | Chr9:33670979..111431608 | 3342 | 774 | 23.16% |
| AR 9.3 | Chr9:131569767..143804017 | 874 | 0 | 0.00% |
| All AR QTL | | 4396 | 774 | 17.61% |
| DPS 3.2 | Chr3:209024681..219411033 | 694 | 0 | 0.00% |
| DPS 8.1 | Chr8:118971709..143492696 | 1462 | 205 | 14.02% |
| DPS 10.1 | Chr10:100252518..128967696 | 1601 | 326 | 20.36% |
| All DPS QTL | | 3757 | 531 | 14.13% |
| GDM/GTN 1.1 | Chr1:3408430..5813154 | 194 | 0 | 0.00% |
| All GDM/GTN QTL | | 194 | 0 | 0.00% |
| PTNP 1.2 | Chr1:43756650..71127215 | 1506 | 0 | 0.00% |
| All PTNP QTL | | 1506 | 0 | 0.00% |
| PTN 2.1 | Chr2:4990077..5565686 | 69 | 0 | 0.00% |
| All PTN QTL | | 69 | 0 | 0.00% |
| PDM 4.1 | Chr4:67143858..237313637 | 8430 | 806 | 9.56% |
| PDM 10.2 | Chr10:147811484..148581023 | 92 | 0 | 0.00% |
| All PDM QTL | | 8522 | 806 | 9.46% |
| SC 1.3 | Chr1:258353240..268813141 | 637 | 0 | 0.00% |
| SC 3.1 | Chr3:10683505..162587460 | 6355 | 936 | 14.73% |
| SC 9.1 | Chr9:22782253..119839322 | 4402 | 820 | 18.63% |
| All SC QTL | | 11394 | 1756 | 15.41% |

**<u>Supplementary Table 4:</u>** <u>AR QTL 1.4 Candidate Genes</u>

**Supplementary Table 4:** Candidate genes within AR QTL 1.4. Columns from left to right indicate Gene Name, Protein Type, and Protein or RNA Upregulation per corresponding tissue type. A green 'Y' indicates upregulation of the corresponding protein or RNA transcript in the corresponding tissue type at young seedling or plantlet stage.

| Gene Name | Protein Type | Confirmed Protein Upregulation | | | | | | Confirmed RNA Upregulation | | | | |
|---|---|---|---|---|---|---|---|---|---|---|---|---|
| | | Root Cortex | Root Elong-ation Zone | Root Meri-stem Zone | Prim-ary Root | Seco-ndary Root | Root Stele | Root Elong-ation Zone | Root Meri-stem Zone | Prim-ary Root | Crown Root Nodes 1 - 4 | Crown Root Nodes 5 - 6 |
| GRMZM2 G129620 | Uncharacterized protein. | Y | Y | Y | Y | | Y | | | | | |
| GRMZM2 G068443 | Uncharacterized protein. | | Y | Y | | | Y | | | | | |
| GRMZM2 G055834 | Uncharacterized protein. | | | | | | | | | Y | Y | |
| GRMZM2 G473906 | Uncharacterized protein. | Y | Y | | Y | Y | Y | Y | Y | Y | | |
| GRMZM2 G477609 | Putative uncharacterized protein. | | | Y | Y | | Y | | | | | |
| GRMZM2 G156848 | Mitochondrial import inner membrane translocase subunit Tim10. | | | | Y | Y | | | | | | |
| GRMZM2 G081105 | Uncharacterized protein. | | | Y | | | | Y | Y | Y | Y | Y |
| GRMZM2 G123876 | Uncharacterized protein. | | | Y | | | | | | | | |
| GRMZM2 G139041 | Uncharacterized protein. | Y | | | Y | | Y | | | | | |
| GRMZM2 G122108 | Uncharacterized protein. | Y | | | | | | | | | | |
| GRMZM5 G840909 | Cytidine/ deoxycytidylate deaminase family protein. | Y | | | Y | | Y | | | | | |
